# Toward quantitative metabarcoding

**DOI:** 10.1101/2022.04.26.489602

**Authors:** Andrew Olaf Shelton, Zachary J. Gold, Alexander J. Jensen, Erin D’Agnese, Elizabeth Andruszkiewicz Allan, Amy Van Cise, Ramón Gallego, Ana Ramón-Laca, Maya Garber-Yonts, Kim Parsons, Ryan P. Kelly

## Abstract

Amplicon-sequence data from environmental DNA (eDNA) and microbiome studies provides important information for ecology, conservation, management, and health. At present, amplicon-sequencing studies – known also as metabarcoding studies, in which the primary data consist of targeted, amplified fragments of DNA sequenced from many taxa in a mixture – struggle to link genetic observations to underlying biology in a quantitative way, but many applications require quantitative information about the taxa or systems under scrutiny. As metabarcoding studies proliferate in ecology following decades of microbial and microbiome work using similar techniques, it becomes more important to develop ways ot make them quantitative to ensure that their conclusions are adequately supported. Here we link previously disparate sets of techniques for making such data quantitative, showing that the underlying PCR mechanism explains observed patterns of amplicon data in a general way. By modeling the process through which amplicon-sequence data arises, rather than transforming the data post-hoc, we show how to estimate the starting DNA proportions from a mixture of many taxa. We illustrate how to calibrate the model using mock communities and apply the approach to simulated data and a series of empirical examples. Our approach opens the door to improve the use of metabarcoding data in a wide range of applications in ecology, public health, and related fields.

## Introduction

Over the past decade, rapid technological advances in the collection and analysis of trace genetic material from sampled environmental media (water (Ficetola et al. 2008, Thomsen et al. 2012), soil (Andersen et al. 2012), feces (Pompanon et al. 2012), or even air (Lynggaard et al. 2022); hereafter environmental DNA [eDNA]) have opened new frontiers for environmental surveillance. Studies using eDNA have focused on diverse topics including monitoring biodiversity (Creer et al. 2016), managing invasive species (Jerde et al. 2013), characterizing diet (Deagle et al. 2013), and supporting fisheries management (Fukaya et al. 2021, Shelton et al. 2022), in habitats from tropical forests (Lopes et al. 2017) to the deep sea (Everett and Park 2018).

These and many other ecological studies use techniques essentially identical to those in microbial ecology, microbiome, and public-health applications, and all share a set of analytical challenges. In most amplicon-based studies (hereafter: “metabarcoding”), a single oligonucleotide primer set targets a region of DNA shared among a taxonomic group of interest (see Taberlet et al. 2012) to be amplified via PCR and subsequently sequenced, with the primer design determining which taxa are likely to be amplified and thus detected. The result is a mixture of DNA sequences from many taxa; the challenge is to determine whether and how the abundances of those sequence-reads correspond to the starting composition of DNA prior to amplification.

There is agreement that metabarcoding data contain information about the taxa present in a sample, and therefore inform estimates of taxonomic richness (reviewed in Taberlet et al. 2018). However, using metabarcoding data to estimate the composition (i.e., taxon-specific proportions) of DNA within a sample is more controversial. Metabarcoding data consists of counts of unique DNA sequences detected by a DNA sequencing machine (e.g. for a given sample we might observe 3 copies of sequence A, 1001 copies of sequence B, etc.). Important bioinformatic decisions bear on how multiple sequences are combined to represent species or genera or higher taxonomic groups (Macé et al. 2022), but the resulting data themselves are straightforward counts of reads associated with particular taxa for each sample. Many uses of these data in an ecological setting share an (often implicit) assumption that the reads emerging from DNA sequencers are an accurate depiction of the sample composition prior to amplification (e.g., Laporte et al. 2021).

There is abundant evidence that the relationship between the true composition of DNA contained in a sample and the reads emerging from the sequencer is far from simple. This fact has been documented in the microbiome and microbial literatures (Gloor et al. 2017, McLaren et al. 2019, Silverman et al. 2021) but less so in the ecological literature (but see e.g., Thomas et al. 2016). The most compelling evidence for this phenomenon comes from the analysis of mock communities in which researchers create a known mixture of DNA from a suite of taxa of interest and compare the relative abundance of the reads from each taxa against the known community. Read counts following amplification and sequencing invariably fail to match – or often, even approximate – the mock-community DNA starting proportions. For example, relative read counts deviate strongly from the mock communities created for freshwater mussels (Coghlan et al. 2021), freshwater invertebrates (Fernández et al. 2018), arthropods (Piñol et al. 2015, Krehenwinkel et al. 2017), freshwater fish (Hänfling et al. 2016, Rivera et al. 2021), marine vertebrates (Port et al. 2016, Andruszkiewicz et al. 2017), fungi (Adams et al. 2013, De Filippis et al. 2017, Palmer et al. 2018), diet studies from a range of organisms (Ford et al. 2016, Thomas et al. 2016, Ando et al. 2020, Tournayre et al. 2020), and microbiome studies (McLaren et al. 2019, Silverman et al. 2021). Beyond their obvious taxonomic diversity, these analyses span a wide range of methodological implementations (i.e. primers and protocols), and are reproducible. Because the differences between expected and observed read-proportions often appear idiosyncratic and species-specific, it is difficult to know how to interpret sequencing reads for quantitative use.

## The Problem

We can illustrate the problem using empirical data from three fish-community datasets: one from British lake communities (Hänfling et al. 2016, Fig. 1A, Fig. 1B), one from Pacific marine communities (this paper; 1C, 1D), and one from fecal diet samples from fish-eating killer whales (*Orcinus orca*, this paper; Fig. 1E, 1F). For each study, we first plot the true proportion of DNA from a mock community of known composition against the estimated proportion of reads detected from each taxon (Figs. 1A, 1C, 1E).

**Figure 1:**
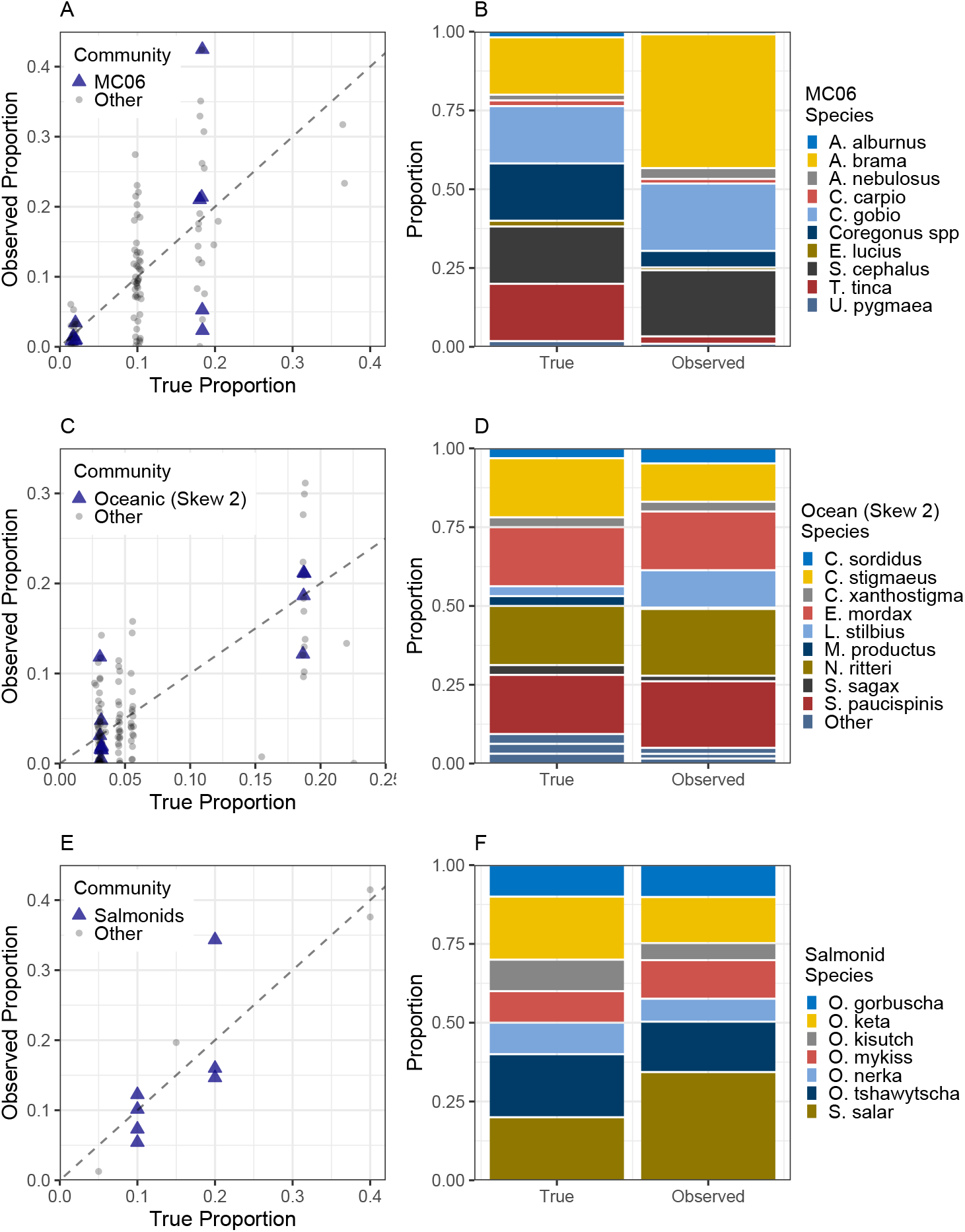
Comparisons between the composition of a known mock community (‘True’) and the estimated proportions derived from raw read counts following sequencing (‘Observed’). Left panels show the relationship for species within mock communities constructed by Hänfling et al. (A; ten mock communities of freshwater fishes; cytochrome b mtDNA), and for this study (C; two mock communities of Pacific Ocean fishes; 12S rRNA; and (E) two mock communities of southern resident killer whale fecal samples; 16S rRNA). Each point (dot or triangle) represents a single species in a mock community. For panel A, each point is a single technical replicate while in panels C and E, each point is the average across three technical replicates. Communities highlighted (triangle symbols) are shown in the stacked bar chart in the right panels (B, D, F).

If the estimated proportions based on sequencing data accurately reflected the original known proportions of the mock community, all points would lie on or very near the 1:1 reference line. While it is reassuring that there is some suggestion of a relationship – larger true proportions are associated with larger observed proportions – points are scattered well above and below the reference line with some points more than double or less than half their true proportions. It is tempting to simply examine Figs. 1A, 1C, and 1E, note that observations are scattered approximately equally above and below the reference line, and conclude that the true proportions are well estimated on average (e.g., Lamb et al. 2019), but to do so would be a mistake. Because these values are proportions of reads associated with each taxon, metabarcoding reads are compositional: the sum of proportions across taxa must equal one, and therefore the points on this graph are not independent. Indeed, if one taxon is above the reference line, one or more other taxa must, by definition, fall below the reference line. Consequently, a positive relationship is very likely for metabarcoding datasets, and, with enough randomly assembled mock communities, the estimated slope will be very near 1. Such a comparison is inappropriate because the lack of independence among the observations renders most standard statistical analyses (e.g., ordinary regression or correlation analyses) inappropriate (see Gloor et al. 2017, Erb et al. 2020).

Examining the true and estimated compositions for a single community (triangles in Figs 1A, 1C, 1E, are shown in stacked barcharts in Figs. 1B, 1D, 1F), it becomes obvious that the true and estimated communities differ substantially. A few taxa match their true abundance closely (see e.g., *O. gorbuscha* in Fig. 1F, *E. mordax* in Fig. 1D) but some taxa are strongly over-represented (*A. brama* in Fig. 1B, *L. stilbius* in Fig. 1D) while others are under-represented (*S. sagax* in Fig. 1D; *O. kisutch* in Fig. 1F).

Two important questions stem from these observations: First, why do these biases in taxonomic composition arise? Second, how do we correct for such biases? We address each in the following sections.

## The Process

While there are several aspects of sequencing data that differ from other types of surveys, PCR amplification and the biases it can introduce into metabarcoding data has repeatedly been identified as the major factor limiting its usefulness (Deagle et al. 2010, 2013, Shelton et al. 2016, Gloor et al. 2017, Krehenwinkel et al. 2017, Kelly et al. 2019, McLaren et al. 2019, Beng and Corlett 2020, Silverman et al. 2021). Thus correcting for PCR-driven biases remains the major challenge for making metabarcoding results reflective of DNA concentrations, although other processes besides amplification can hinder the reconstruction of the relationship between metabarcoding data and taxonomic abundance (e.g., variability in DNA deposition and persistence, copy number variation, biases in taxonomic assignment, Gohl et al. 2016, McLaren et al. 2019, Beng and Corlett 2020). Understanding the mechanisms by which metabarcoding data are produced is therefore essential for reliably understanding PCR-based data.

We adopt a structure for understanding PCR-driven biases that has been used previously (Kelly et al. 2019, McLaren et al. 2019, Silverman et al. 2021) and allows PCR-driven variation to be a function of variation among taxa in PCR amplification efficiency. Specifically, for any taxon *i*, the number of sequence-reads produced during a PCR reaction are governed by an efficiency parameter *a*_*i*_, which is characteristic of the interaction between the particular primer set and each taxon (or unique sequence variant) being amplified. Thus, for any given taxon, we expect the number of reads to be directly related to the efficiency of amplification and the starting concentration of DNA template. Let *A*_*i*_ be the expected number of sequence reads after PCR, *c*_*i*_ be the true number of DNA copies in the reaction attributable to taxon *i, a*_*i*_ be the amplification efficiency (bounded on (0, 1)), and *N*_*PCR*_ is the number of PCR cycles used in the reaction.

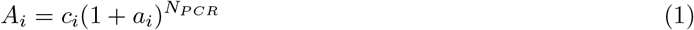

Note that this equation only applies during the exponential phase of PCR, before reagents have been exhausted and the amplification process has stopped, a valid assumption for most eDNA applications which start with very low DNA concentrations. If we could perfectly observe the DNA molecules, the above equation alone would be sufficient to understand the value of interest, *c*_*i*_. Unfortunately PCR and sequencing technology does not allow for such direct observation. For any useful primer set, *a*_*i*_ is typically not close to 0 (a value of 0 would indicate no amplification during PCR) and *N*_*PCR*_ is large (*>* 30), so the number of sequence reads expected for any taxon with *c*_*i*_ *>* 0 is very large (e.g., with *c*_*i*_ = 2, *a*_*i*_ = 0.75, and *N*_*PCR*_ = 36, *A*_*i*_ = 1.12 ×10^9^). Given the simultaneous amplification of many taxa, 10^10^ or more DNA copies are typically produced.

DNA sequencing instruments report only a small fraction of the total amplicons following amplification (often on the order of 10^6^ to 10^7^ reads per sample, depending upon the sequencing instrument, indexing decisions, and so on). Thus only a small fraction of the total generated amplicons is actually observed. Assuming that the sequencing mechanism reports an unbiased subsample of amplicons, we can think of the observed reads for each taxon (*Y*_*i*_) as proportional to the true amplicon abundance: *Y*_*i*_ ∝ *A*_*i*_. This sampling changes what in eq. 1 appears to be a single-taxon process – each taxon being amplified independently – into a multi-taxon, compositional process; the number of amplicons observed for taxon *i* will depend both upon the amplicons produced for taxon *i* = 1 and the amplicons from taxa *i* = 2, 3, …, *I* in the same reaction.

The consequences of (1) among-taxon variation in amplification rate, and (2) the compositional nature of the resulting data for inferences about ecological communities are profound. We provide two graphical examples of the potential effects of amplification variability on inferences of ecological communities consisting of three hypothetical taxa (Fig. 2). Ecologists are interested in quantifying the DNA present before any PCR amplification (shown as PCR cycle 0 in Fig. 2), but if taxa differ in their amplification efficiency (*a*_*i*_), the relative abundance of amplicons can shift dramatically over the course of PCR amplification such that when the reads are observed following sequencing (filled points), the composition of the observed sequences differ substantially from the composition of starting DNA molecules. Note that the relative abundances and even the rank order of abundances can change between the initial and final PCR cycle (0 and 30, respectively; Fig. 2). Furthermore, it is vital to understand that this is a multivariate process; the observed relative abundance of any one taxon is dependent upon the other taxa co-occurring with it in the sample. Compare the fate of taxon “A” in the two different communities depicted in Fig. 2. In both panels, taxon “A” initially comprises one-sixth (≈ 17%) of the community but in the first community it makes up more than 60% of amplicons after PCR amplification (Fig. 2A); however, in the second community it declines slightly to ≈ 16% of amplicons after PCR amplification (Fig. 2B). This is not a result of anything fundamental changing about taxon “A” but rather the fact that a taxon with a higher amplification efficiency (taxon “D”) comprises half the community in Fig. 2B while it is replaced by a taxon with lower efficiency (taxon “C”) in the community in Fig. 2A.

**Figure 2:**
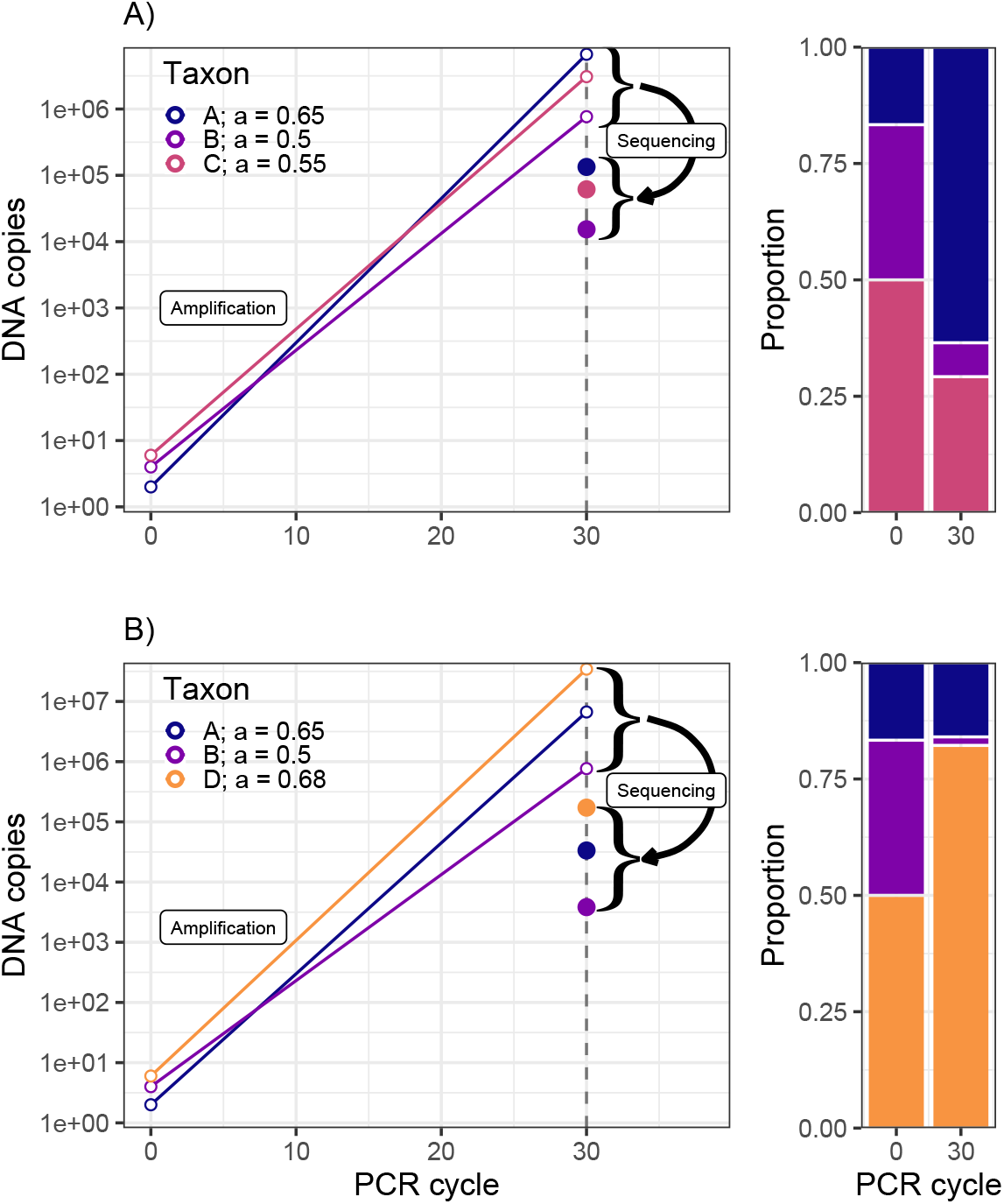
Two examples of simulated changes in communities due to PCR bias. Each shows communities of three taxa with variation in amplification efficiency. Left panels illustrate the PCR process and denote the true (but unobserved) number of DNA copies present (open circles) and as well as observed counts following DNA sequencing (filled circles). Slopes of lines are determined by the taxon-specific amplification efficiencies (*a*_*i*_). Right panels compare the true starting-community DNA composition at PCR cycle 0 and the observed community composition at PCR cycle 30. Taxa A and B are shared across communities, with identical starting proportions in each, but radically different proportions following PCR.

While Fig. 2 presents stylized examples, it makes four important points. First, varying amplification efficiencies among taxa have the potential to dramatically affect the amplicon counts observed after sequencing. Second, the patterns of bias produced by allowing for amplification variation qualitatively match the patterns observed in the empirical analysis of mock communities (Fig. 1). Third, in the presence of varying amplification efficiencies, it is impossible to determine the initial composition of a sample simply by observing the relative abundance of amplicons associated with each taxon after sequencing; many possible combinations of parameter values for starting proportion and amplification efficiency would yield the same observed proportions post-PCR. Finally, interpreting amplicon counts cannot be done on a taxon-by-taxon basis, but must be done in a multivariate context. We can therefore quickly identify and disregard approaches that will clearly not resolve the problem. Specifically, simple data transformations – including logarithms, roots, or any other monotonic transformation – will not succeed in correcting for biases introduced by amplification variation among taxa (e.g., Kelly et al. 2019).

## Methods

Various techniques have been proposed to reconcile the differences between the amplicon data we observe and the abundances of the underlying organism-specific DNA concentrations in the environment. These range from process-based statistical approaches solidly grounded in theory (McLaren et al. 2019, Silverman et al. 2021), to laboratory methods such as tagging molecules individually prior to amplification to distinguish replicate amplicons from unique template molecules after sequencing (e.g., qSeq, Hoshino and Inagaki 2017, Hoshino et al. 2021, and other Molecular ID tags (MIDs)) to various post-hoc transformations and corrections (Thomas et al. 2016, Krehenwinkel et al. 2017, Kelly et al. 2019).

Here we treat observed sequence reads as arising from the mechanics of the PCR reaction and subsequent DNA sequencing, developing a statistical model grounded in the PCR process itself. We set out a quantitative model for amplicon data to account for the effects of amplification bias, estimating the proportion of each taxon’s DNA in the original PCR template (i.e., community composition of our samples) prior to PCR and sequencing. The key to doing so is estimating taxon-specific amplification-efficiency parameters. We use multinomial logistic regression to account for PCR bias, and make clear the general applicability of these models to all kinds of amplicon-based studies. We emphasize that the models we use are not unique – other researchers have provided closely related approaches in the medical and microbiome literature (e.g., McLaren et al. 2019, Silverman et al. 2021) – but these techniques are underused and underappreciated in the ecological literature. We also note that there are some similarities between existing correction procedures (Thomas et al. 2016, Krehenwinkel et al. 2017) and parts of our approach.

We summarize the primary statistical model in the main text and present extensions in Supplement A. After the model description, we highlight the calibration requirements for these models and emphasize how adding a few steps to molecular data-collection protocols can greatly improve their value for ecological inference. Given the potential for uncorrected data to lead to potentially large errors in ecological inference, we aim to make a complex statistical model approachable to most practitioners.

### Compositional Models for Amplicon Data

The sampling process associated with DNA sequencing severs the link between the absolute abundance of the initial DNA copy count or concentration (*c*_*i*_; see eq. 1 and 2) and the count of DNA sequences observed (the *Y*_*i*_). Gloor et al. (2017) present illuminating examples illustrating the challenges of compositional data. Specifically, we can write the ratio of observed sequences for taxa *i* and *j*, 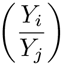 as

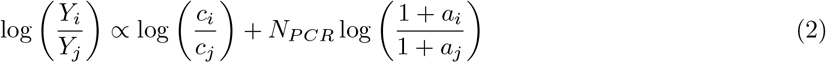

a function of the initial ratio of taxa *i* and 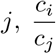, modified by the product of the ratio of amplification efficiencies for the two taxa, 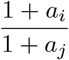, and the number of PCR cycles. Note that in eq. 2 there are many possible values that yield the same ratios (e.g. the pair {*c*_*i*_ = 2, *c*_*j*_ = 1} produce the same ratio as the pair {*c*_*i*_ = 4, *c*_*j*_ = 2}; 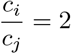), emphasizing the loss of information about absolute scale when dealing with compositional data and ratios.

To solve this scaling problem, we can arbitrarily define one taxon to be a reference taxon (*R*) and define a new set of parameters relative to this reference taxon. Let *β*_*R*_ = 0 be the log-abundance of the reference taxon in the initial sample and *β*_*i*_ be the abundance of taxon *i* relative to the reference (*β*_*i*_ *>* 0 indicates taxon *i* is more abundant than the reference, *β*_*i*_ *<* 0 the opposite). Similarly, let *α*_*i*_ be the log-efficiency relative to the reference taxon (*α*_*R*_ = 0), then *ν*_*i*_ is the log-abundance of taxon *i* relative to the reference taxon after sequencing.

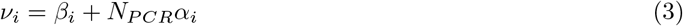

By definition, *ν*_*R*_ = 0, and the choice of reference taxon is arbitrary, but not unimportant. In this formulation, *ν*_*i*_ is a linear function having slope *α*_*i*_ and intercept *β*_*i*_; the intercept determines the proportion of DNA for taxon *i* present in the sample before PCR, and the equation defines the proportional abundance for any number of focal taxa.

We acknowledge the stochastic processes that contribute to the observed read counts beyond the deterministic skeleton presented in eq. 3, and we can model this stochasticity using a multinomial likelihood (Egozcue et al. 2020, Silverman et al. 2021). A full model for observed counts for all *I* taxa is

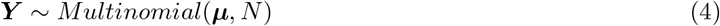

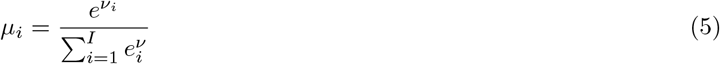

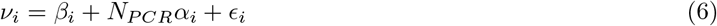

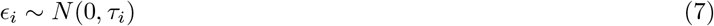

where bold text indicates vectors and *N* is the observed total number of sequences in the sample. Equation 5 is the softmax transformation which produces proportions for each taxon (*µ*_*i*_) from the ratios of abundance. The parameter allows for over-dispersion in the counts beyond the variability provided by the multinomial distribution, capturing the substantial variance among technical replicates often observed in metabarcoding data. As will be seen below, *ϵ* can be important because read-counts may vary substantially across replicate PCR reactions, but this may not be estimable for some metabarcoding datasets, which commonly lack technical replication. The model described in eq. 4 is known as a multinomial logistic regression model, and defining log-ratios relative to a reference taxon is known as the additive log-ratio transform (ALR, Aitchison 1986).

Above we have written a simple scalar-valued form to improve readability, but the regression component quickly generalizes and can take on more complicated structures from the diverse world of generalized linear models (e.g., Silverman et al. 2021). Ecologists are rarely interested in a model for a single sample as written in eq. 4 but in a model for many samples taken across space (e.g. latitude, habitats) and/or time (seasons, months, years) and the model generalizes to accommodate such questions. For example, in the context of diet data it may be valuable to add terms describing measured covariates (e.g. temperature) or factors such as season, and the stochastic term can be modified to allow for additional covariance structure among samples. We present a general model in Supplement 1 to make such modifications explicit. Similarly, it may be desirable to use transformations other than the ALR depending on the application (e.g., centered log-ratio transform (CLR) or isometric log-ratio transform, Pawlowsky-Glahn and Egozcue 2016, Silverman et al. 2021).

Most metabarcoding datasets include a single sequencing replicate for each sample collected after a single PCR amplification (corresponding to a single *N*_*PCR*_) and thus have only *I* observations (one sequence count for each taxon). The model above requires at minimum *I* − 1 *β* parameters and *I* − 1 *α* parameters. Thus with standard metabarcoding data, there are more parameters than data points and this model cannot be estimated. In the next sections we discuss how to integrate other data sources to calibrate the model, make the parameters identifiable, and allow researchers to generally correct for amplification bias.

### Calibration methods

There are at least four general strategies that can provide the information necessary to estimate amplification biases: 1) create mock communities of known DNA template composition and conduct PCR amplification and sequence this known community; 2) use samples of unknown composition, but vary the number of PCR cycles among technical replicates and subsequently sequence all replicates; 3) model amplicon results alongside another independent set of observations of the same community; 4) attach unique molecular identifiers to source molecules prior to amplification and analyze unique identifiers post-PCR. We focus on the first approach here because we believe it to be broadly applicable and we have had success implementing it in practice. Silverman et al. (2021) provide an investigation using variable PCR cycles applied to microbiome data. Gold et al. (In Review) provides a substantial treatment of the third approach – there, with amplicons and visual counts of larval fish communities in ethanol-preserved jars. Hoshino and colleagues review the fourth approach (qSeq, Hoshino and Inagaki 2017, Hoshino et al. 2021).

To provide intuition about how these methods allow us to estimate amplification rates, we present two simple graphical illustrations for approaches using mock communities and variable PCR (Fig. 3). For both approaches, the amplification efficiency for each taxon is directly related to the slope of the line in Fig. 3 (see also eq. 3).

**Figure 3:**
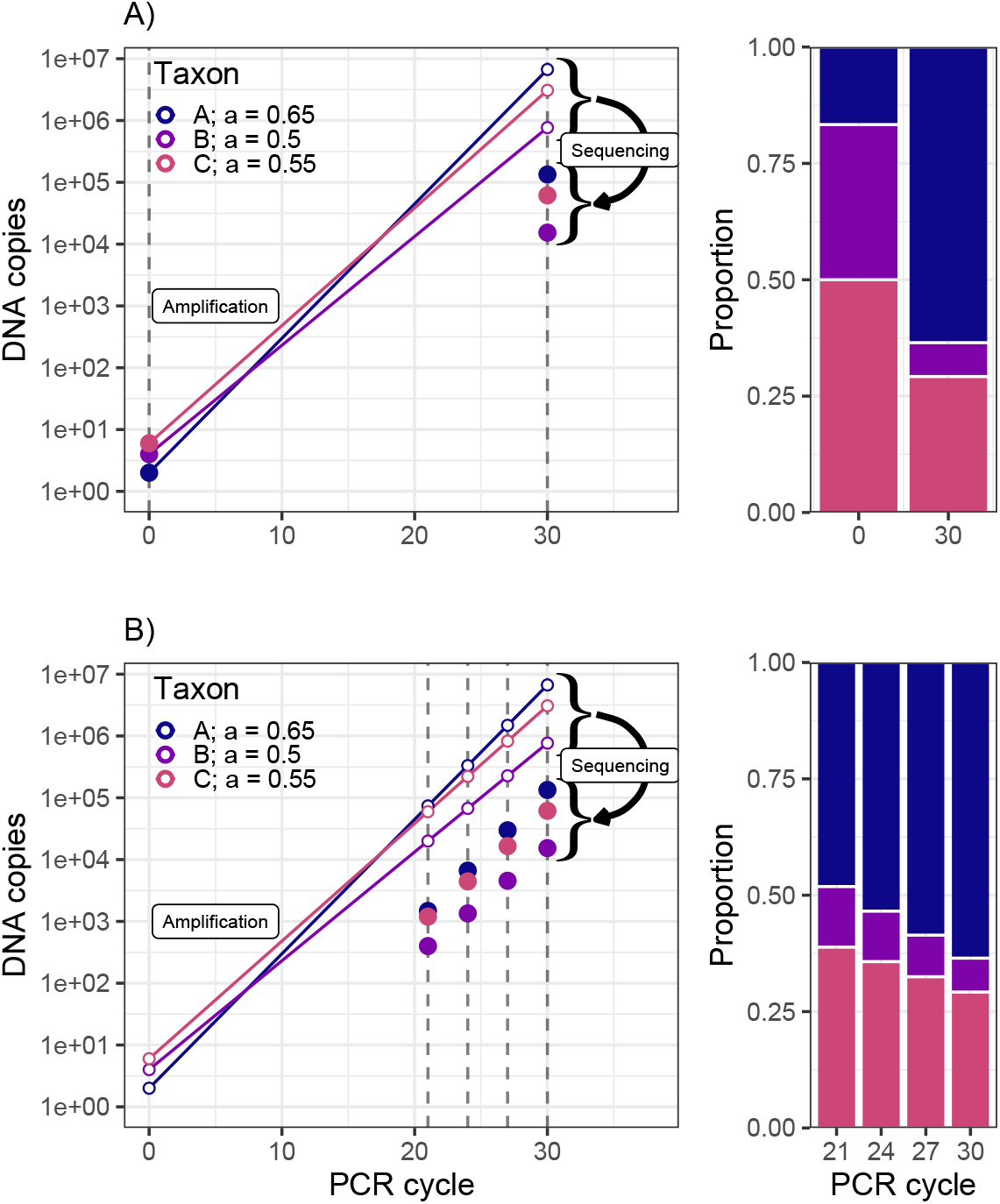
Two schematic illustrations of techniques to enable estimation of amplification-bias among taxa. In all panels, filled circles indicate known values or observations and empty circles indicate unknown states. The mock community approach (A) provides known initial taxon composition (PCR cycle 0) that can be compared to the composition of sequences observed at PCR cycle 30 to estimate relative amplification rates. The variable PCR cycle calibration (B) uses a sample with unknown community composition at PCR cycle 0, but observes the community of amplicons multiple points in the PCR process, sampling at different numbers of amplification cycles. With this method, the change in relative community composition across PCR cycles provides information about relative amplification rates. The right panels of both A and B show the observed composition of the samples after a given number of PCR cycles.

### Mock communities

To calibrate using a mock community, we create the starting community of DNA, generally from vouchered tissue-extractions. We therefore we know the proportions of each taxon before PCR (cycle 0) and we observe the amplicons following sequencing (Fig. 3A). Thus we have an estimate of the initial and final community composition (filled points in Fig. 3A), allowing us to estimate the relative amplification efficiency (***α***, relative slope parameters) for each taxon. We can then apply these estimates of ***α*** to samples of unknown community composition, yielding an estimate of the parameters of ecological interest, ***β*** (intercept parameters in eq. 4; Fig. 3A). In practice it makes sense to use joint models that simultaneously incorporate observations from mock communities alongside samples of unknown composition. We adopt this technique in our simulations and empirical applications below.

### Variable PCR cycles

While mock communities offer a reasonable approach to estimating amplification biases, they have several drawbacks. Most obviously, they require constructing an appropriate mock community for a given application. When there are large numbers of taxa of interest or source DNA from important taxa are unavailable, creating a mock community of known composition may be difficult or impossible – as is often the case in microbial community studies, for example. In such cases, it is possible to modify the PCR protocol for technical replicates to bracket a range of PCR cycles and observe the amplicons at each end point (Fig. 3B). Importantly this variable PCR cycle calibration can be applied to samples of unknown composition. In regression terms, the intercept parameters (***β***) are unknown in the variable PCR cycle approach, but the change in relative composition across a range of PCR cycles enables estimation of the relative amplification of different taxa (Fig. 3B), ***α***, and in combination with observed proportions of sequences after PCR, yields estimates of starting proportions ***β***. While this approach has been used to good effect with significant sampling effort (Silverman et al. 2021), we have not had success using it in practice. We include it here to illustrate its appealing logic.

## Applications

The quantitative model described above is only valuable if it can be useful in practice. We developed code to implement the model outlined in eq. 4 in *Stan* via the **R** language (*Rstan*) and then tested it against several data sets, both simulated and empirical. First, we provide simulations to illustrate the model’s ability to recover known parameters. We then apply the mock-community calibration method to several empirical examples to estimate community composition. Each empirical example is drawn from a different ecological context; these include British lake fishes (unreplicated PCR reactions from mock communities, *cytochrome b mtDNA*; data from (Hänfling et al. 2016)), Pacific Ocean fishes (replicated PCR reactions from mock communities, *12S rRNA*; original data), and diet data derived from the gut contents of southern resident killer whales (SRKW; replicated PCR reactions, *16S rRNA*; original data). We present details of model estimation and prior distributions for the model in Supplement A. Because we focus here on eukaryotic communities, we include the above examples in the main text, but for completeness we also show an analysis of human bacterial microbiomes (replicated PCR reactions, *16S rRNA*; data from (Gohl et al. 2016)) in Supplement 1; *16S* bacterial data were also shown in Silverman et al. (2021) and McLaren et al. (2019). All data and code used in the applications are provided in the online supplement.

### Simulation testing

We first generated sets of simulated amplicons arising out of linked PCR and sequencing processes. We simulated observations of amplicons for 25 species in 10 biological samples in a PCR reaction, observing an expected total of 10^6^ reads per replicate for 3 independent technical replicates. Each simulated taxon was assigned a known, randomly selected amplification efficiency, *a*_*i*_ and we simulated a PCR protocol with *N*_*PCR*_ = 35 cycles. Proportional contributions for each taxon were drawn from a symmetric Dirichlet distribution with parameter *α*_*D*_ for each combination was 0.2 to achieve high variability among samples. To simulate amplification, we followed eq. 1 but allowed for overdispersion in counts with a negative binomial distribution, and then specified compositional sampling using a multinomial distribution,

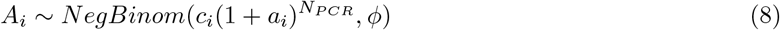

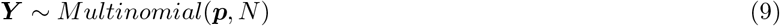

where the negative binomal is parameterized in terms of its mean and overdispersion and 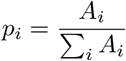.We evaluated model performance for six scenarios of varying amplification efficiency among taxa and overdispersion among technical replicates. Specifically, we evaluated amplification efficiencies with low, moderate, and high variability among taxa by drawing *a*_*i*_ from a beta distribution (*a*_*i*_ ∼ *Beta*(*δ, δ*)) with *δ*= 50, 10, or 2 (i.e., low, moderate, and high amplification variability, respectively). For each level of amplification variability, we also simulated low and high variability among technical replicates (*ϕ*=10, 2, respectively). For the mock community calibration, we simulated four communities with three technical replicates apiece. One mock community included all simulated taxa at equal proportions, and the remaining three communities each included approximately half of the taxa at equal proportions (i.e., include taxa 1 to 12, taxa 6 to 18, and taxa 13 to 25 in separate communities). Sample sizes from the mock community training data represented 40% of the out-of-sample sample size, which included data from 10 biological samples. We summarized model fit relative to the known starting community composition using Aitchinson distance [AD; Aitchison (1982)], as is appropriate for compositional data (Gloor et al. 2017). For AD, smaller values indicate more similarity between two compositions.

## Empirical Examples

### British Lakes

Hänfling et al. (2016) sampled eDNA from a series of freshwater lakes in northern England, and alongside these environmental samples, reported data from 10 cytochrome-b mock communities having known starting compositions and consisting of partially-overlapping mixes of 21 species drawn from the target lake-fish communities. These samples were amplified in triplicate but the three reactions were pooled and each mock community was sequenced once on an Illumina MiSeq. See Hänfling et al. (2016) for analytical and bioinformatic details of the original dataset; the authors provided read-counts, compositions for mock communities, and analytical code as part of the publication’s supplementary material (https://github.com/HullUni-bioinformatics/Haenfling_et_al_2016).

We took species-specific starting DNA concentrations and resulting read-numbers from the authors’ Table S5, and consistent with our method of subsetting data to the relevant target group, omitted any reads from species not included in the mock community preparations (i.e., for which starting concentrations were zero; Table C1). We note that this approach is not equivalent to simply ignoring contamination by selectively omitting data (see Discussion). Instead, it focuses the analysis on the subgroup of interest, and results in estimated proportions for species in that subgroup, rather than for the sequencing run as a whole. For purposes of model-fitting, we treated all PCR reactions as having 30 amplification cycles (Appendix 6, Hänfling et al. 2016).

For model-fitting and cross-validation we created a version of the metabarcoding model with no overdispersion term (i.e. fix *ϵ* = 0 in eq. 4), because in the absence of technical replicates it is impossible to estimate overdispersion and so the read counts are assumed to follow a multinomial distribution. We fit two models, the model described above and a model which assumes equivalent amplification among all species (i.e. fix *α* = 0 and *ϵ* = 0 in eq. 4). For each model we then used the odd numbered mock communities (1,3,5,7,9) as samples with known starting concentration, and even communities (2,4,6,8,10) as unknowns to be estimated. Because we know the starting composition of each of those communities, we used this second set for external cross-validation. We then did the reciprocal cross-validation, using even numbered communities as known and odd numbered communities as unknown, giving us a complete set of 10 cross-validated, out-of-sample estimates. We compare posterior estimates of the community compositions relative to the known starting community composition of the mock community using AD (Aitchison 1982).

### Pacific Ocean Fishes

To analyze a larger suite of species and to estimate overdispersion among technical replicates, we extracted DNA from voucher tissue samples from the Scripps Institute of Oceanography Marine Vertebrate Collection, generating template communities of temperate fish communities from the Northeast Pacific (see detailed methodologies for our entire protocol in Supplement C). We generated two species pools: “North,” comprised of 22 fish species found in coastal Alaskan waters, and “Ocean,” containing 18 species representative of fish found in the California current system (Table C2). Six species were present in both pools. For each species pool we then constructed 3 mock communities with varying composition. In one community, all species were equally abundant (“even” community). In the other two, we selected 12 species, four species each comprised 18.75% of the community and the remaining 8 species comprised 3.125% (“skew 1” and “skew 2” communities; Table C2). We amplified each mock community in separate triplicate reactions using the MiFish Universal Teleost 12S primer set (Miya et al. 2015) at 39 PCR cycles. Supplement C provides further information about assembly of the mock communities and about bioinformatic processing.

As with the British lakes data, we divide our data into two groups, a set used for estimating amplification parameters and a set held out for out-of-sample cross-validation. We used two communities (Ocean even, North even) as mock communities with known composition and the remaining four (Ocean skew 1, Ocean skew 2, North skew 1, North skew 2) to calculate out-of-sample predictive accuracy. Thus, we used a one-third of our mock communities to predict the remaining two-thirds. Each community had three technical replicates and therefore we used the full model as described in eq. 4. We also fit a second model assuming equivalent among-species variation in amplification (fix *α* = 0 in eq. 4) and used Aitichson distance to compare the predicted vs. true community composition for each model.

### Killer Whale Diet

To understand the impact of amplification bias on diet estimation, we examined data generated from a small subset of fecal samples collected from SRKW. SRKW are fish-eating whales that live primarily along the coast of Washington state, USA and British Columbia, Canada, and are listed as Endangered under the Endangered Species Act. Understanding SRKW diet is important for understanding prey availability and supporting the recovery of their population (Hanson et al. 2021). We combine mock communities for common SRKW prey items with a small number of field collected fecal samples to understand how amplification bias may change estimates of diet composition.

We generated two mock communities representative of SRKW prey species from genomic DNA extracted from individual, vouchered fish-fin or muscle samples (Ford et al. 2016). Whole genomic DNA was normalized to a concentration of 0.5ng/ul using a qPCR SYBR assay of a fragment of the 16S SSU rRNA gene, before being combined in known proportions into two mock communities comprising 10 species (7 salmonid species and three other species; Table C4).

We included eight fecal samples of unknown species composition (‘field samples’) to analyze alongside the mock communities. Samples were collected during the month of September in 2017 (n = 6), 2018 (n = 1), and 2021 (n = 1). We amplified DNA extracts with primers targeting the 16S rDNA region (Ford et al. 2016) for both mock communities and field samples, using a 32-cycle PCR with two technical replicates for each field sample and four technical replicates for each mock community. Full laboratory and bioinformatic protocols are described fully in Supplement C, as is information for incidental take permits under the U.S. Endangered Species Act and the U.S. Marine Mammal Protection Act.

Unlike in the British Lakes and Pacific fish examples, we do not have additional, known SRKW diet communities with which to make out-of-sample estimates of model accuracy. Instead, we estimate two models (one with amplification variability (eq. 4) and one without amplification variability (eq. 4 but with *α* = 0)) to illustrate how including amplification biases modifies estimated diet composition field samples. For analysis, we include only the ten prey species included in the mock community and excluded other rare species arising in the sequenced samples; in total other species never comprised more than 0.65% of reads in any sample. We present predicted compositions for individual samples as well as used the posterior estimates for each individual fecal sample to derive an average diet composition from these eight samples.

## Results

### Simulations

Our simulations suggest that calibration with mock communities effectively corrects for amplification bias (Fig. 4). With limited variability among technical replicates, calibration recovers taxon starting-proportions accurately and precisely when amplification efficiencies have low to moderate variability among taxa (Fig. 4A,C). The mean posterior estimates are unbiased and almost perfectly correlated with the true proportions, and credible intervals are small; model estimates uniformly better approximate true proportions than do observed proportions (mean AD among samples and replicates=7.50, 8.93 for low- and moderate variability in amplification efficiency, respectively, relative to a model based on observed proportions alone; mean AD=10.49, 13.93). With highly variable amplification efficiencies among taxa – and in particular when many taxa amplify poorly – estimated species proportions are still strongly correlated with the true values, but estimates become far less precise (compare Fig. 4A with Fig. 4E). ADs with high amplification variability are accordingly higher (mean AD = 15.22 for estimates based on mock communities and 21.65 for observed reads alone).

**Figure 4:**
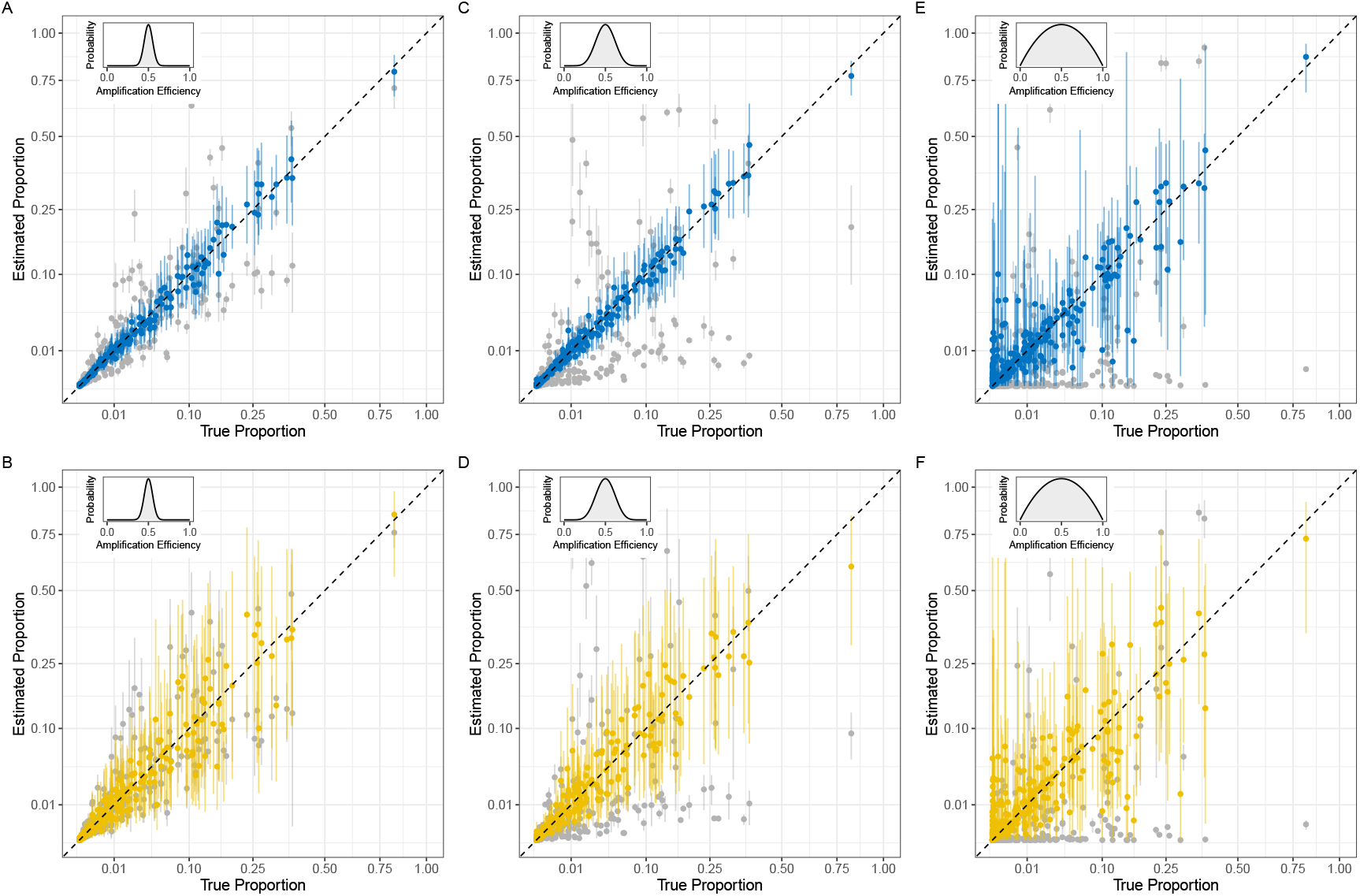
Observed data (grey points) and posterior model estimates (colored points with 95% CI) for simulations of 25 species from 10 sample populations. Panels A, C, and E (in blue) show estimates for datasets with a moderate level of variation among triplicate technical replicates (Negative Binomial *ϕ* = 10 ; low overdispersion). Panels B, D, and F (in yellow) show estimates for a greater level of variation among replicates (Negative Binomial *ϕ* = 2 ; high overdispersion). Inset plots show the probability distribution from which primer amplification-efficiencies were drawn for the data shown in each panel; the variation among amplification efficiencies increases from left to right.

Similarly, increasing the variability in read counts among technical replicates results in increased uncertainty in estimated species compositions (Fig. 4B,D,F). Model performance was consistently worse with greater variability in amplification efficiencies but accounting for amplification bias improved estimates of species composition (mean AD = 8.02, 9.14, 16.08 with low, moderate, and high amplification variability, respectively, from mock communities and mean AD = 11.27, 14.56, 21.62 for observed reads alone).

### Empirical Data

#### British Lakes

We estimated substantial variation in the relative amplification efficiencies (*α*_*i*_) of lake fish species, which varied over roughly 0.3 units (Fig. 5D; we used *Abramis brama* (common bream) as the reference species; *α*_*R*_ = 0). Estimated amplification efficiencies derived from using the odd-versus even-numbered communities for calibration were quite similar, suggesting amplification efficiency derived from different mock communities are consistent. In general, lower-efficiency species have greater uncertainty because they are observed rarely or are absent entirely in the observed sequences (Fig. 5D).

**Figure 5:**
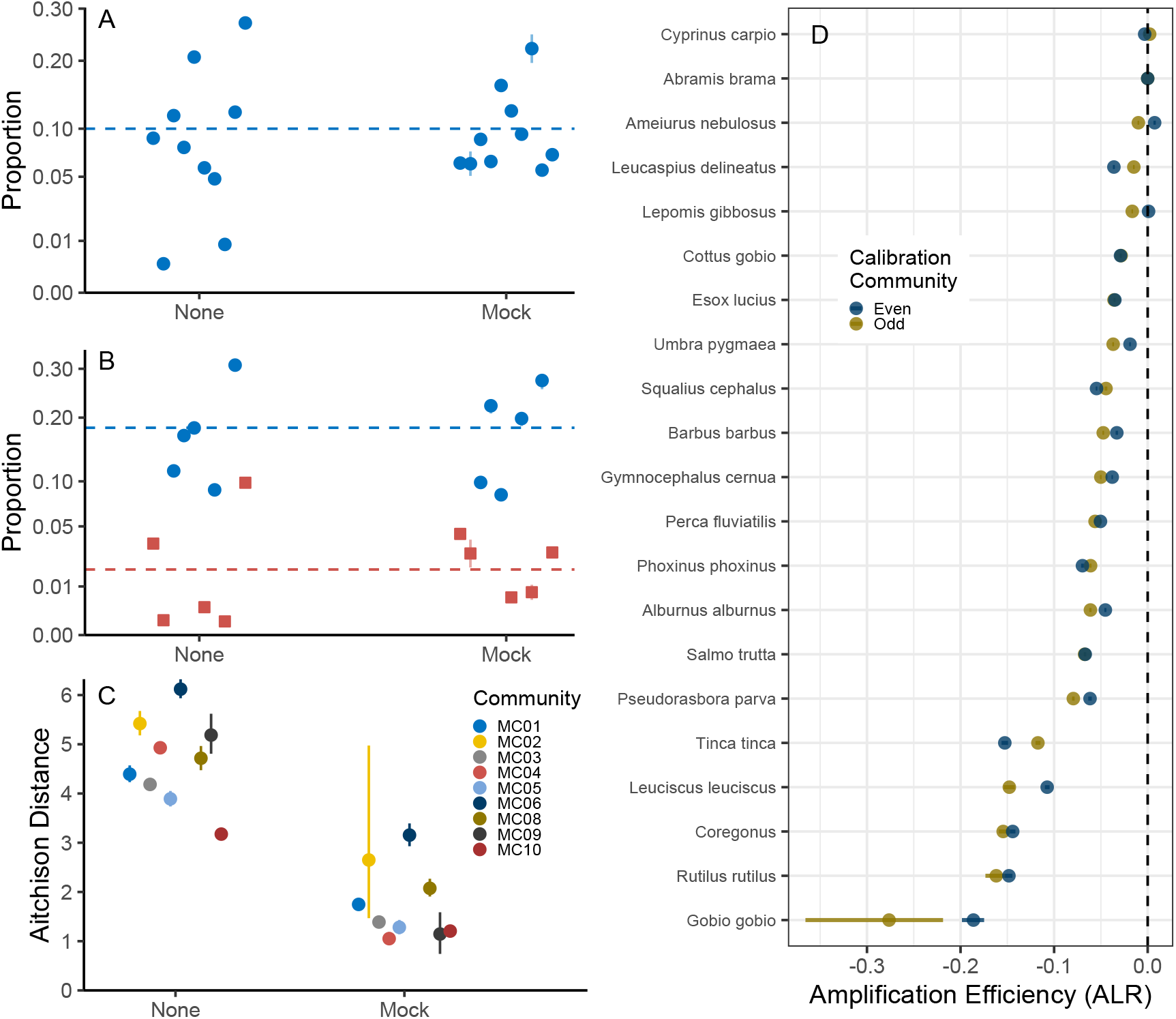
Comparison of calibration methods for British Lakes fish communites. A: Posterior mean estimates (95% CI) of species composition for the ten species in the mock community MC03 without estimated amplification variability (‘None’) or with amplification variability estimated using a mock community (‘Mock’). Dashed line shows the true composition for each species. B: Posterior mean estimates (95% CI) of species composition for the ten species in the mock community, MC08. Dashed blue line shows the true composition for the species identified with a circle. Dashed red line shows the true composition for the species identified with a square. Otherwise as in A. C: The similarity between the true composition and the estimated composition as measured by Aitichson distance. Posterior mean and 95% CI shown. Smaller values indicate greater similarity. D: Posterior mean estimates (95% CI) of relative amplification efficiency derived from two two calibration sets (Odd-numbered communities used as mock community or Even-numbered communities used as mock community). *Abramis brama* is the reference species (*α* = 0) for both communities.

Model estimates of community composition more closely approximated the true starting concentrations (relative to a null model assuming a constant *α* across species) in all 10 communities in cross-validated out-of-sample predictions (Fig. 5). We illustrate this improvement for two example communities (Fig. 5A,B), as well as in the summarized Aitchison distance for all communities (Fig. 5C; note that community MC07 had much larger Aitchison distance; see Fig. A2, A3).

Despite the improved fit relative to the true composition, the credible intervals for all parameters were unreasonably small and only rarely did the credible interval for estimated proportions include the true composition. For example, no species has credible intervals that include the true proportion in Figs. 5A and 5B, indicating overconfidence in estimates of composition as well as amplification efficiency (Fig. 5D). This is largely a result of having a single data point for each species in each community, and the necessary assumption of multinomial sampling variability. As we show directly below, technical replication can resolve this issue.

#### Pacific Ocean Fish

For mock communities of 34 Pacific Ocean fishes, we estimated substantial variation among species in the amplification efficiency with the 12*S* MiFish primer (Fig. 6E). Note that rather than presenting direct estimates of *α*_*i*_ as in Fig. 5, we transformed our amplification using the centered log-ratio transform (CLR; 0 in Fig. 6, which uses the geometric mean among species as 0 rather than the reference species as 0). This allows for consideration of taxon-specific amplification efficiencies relative to the average efficiency in the community.

**Figure 6:**
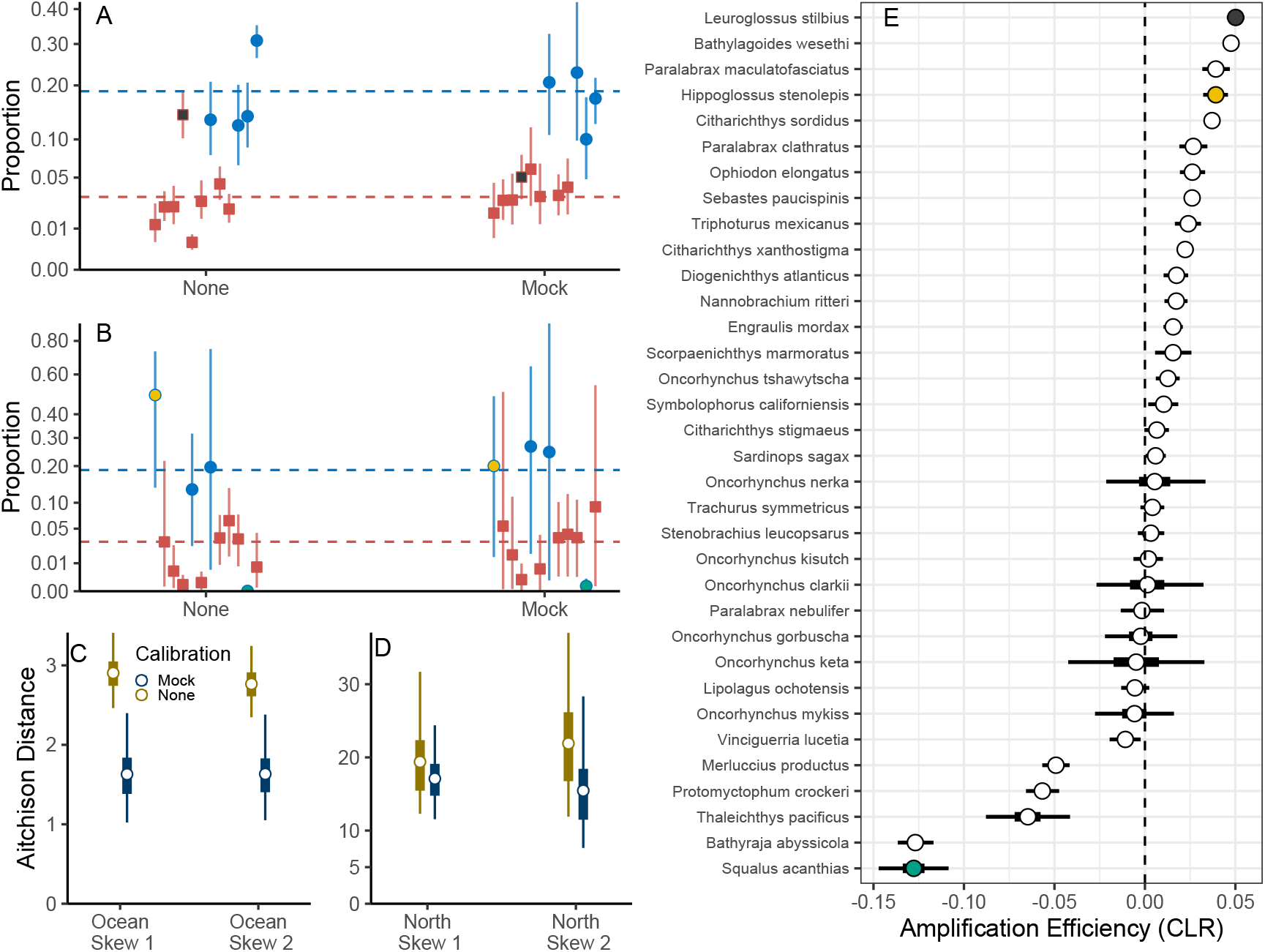
Comparison of calibration methods for Pacific ocean fish communites. A: Posterior mean estimates (95% CI) of species composition for the 12 species in the mock community Ocean skew 1 without estimated amplification variability (‘None’) or with amplification variability estimated using a mock community (‘Mock’). Dashed blue line shows the true composition for the species identified with a circle. Dashed red line shows the true composition for the species identified with a square. Colors correspond to species noted in panel E. B: Posterior mean estimates (95% CI) of species composition for the 12 species in the mock community, North skew 2. Otherwise as in A. C and D: The similarity between the true composition and the estimated composition as measured by Aitichson distance for the four out-of-sample predicted communities. Posterior mean, interquartile range, and 95% CI shown. Smaller values indicate greater similarity. E: Posterior mean estimates (95% CI) of relative amplification efficiency after centered log-ratio transformation. Dashed line indicates the geometric mean amplification efficiency among species. Colors correspond to the colors in panel A and B. *Citharichthys sordidus* was used as the reference species.

In line with simulation and the British lake data, models that account for amplification variability produced estimates of species composition more similar to the true, underlying composition than models that did not account for amplification variability for all communities examined (Fig. 6A–D). However, in contrast to the British lakes data, credible intervals for each species included the true species composition for 11 of 12 species in the Ocean skew 1 community (Fig. 6A) and 10 of 12 species in the North skew 2 community (Fig. 6B). Without calibration, the credible interval included the true proportion in 8 of 12 species in both Ocean skew 1 and North skew 2 (Fig. 6A,B). Larger credible intervals better reflect the variability common in metabarcoding datasets, and here are a direct result of sequencing multiple technical replicates and allowing the model to estimate overdispersion (*τ*_*i*_ in eq. 4). Estimates of *τ*_*i*_ range from 0.25 to 3.9 (posterior mean) among species indicating substantial variability relative to the multinomial distribution.

We highlight the behavior of a few key species with colors in Fig. 6 to emphasize the effect of amplification efficiency on model predictions. For species with higher-than-average amplification efficiency (large *α*_*i*_), the posterior estimates after calibration closely approximate the true species proportions even when the uncalibrated observations of those species are particularly misleading (see *Leuroglossus stilbius* in Fig. 6A,E and *Hippoglossus stenolepis* in Fig. 6B,E). Conversely, species that amplify poorly offer little information on which to base model fits, and so the posterior estimates will have little relationship to their true values (*Squalus acanthias*, Fig. 6B,E). Despite comprising 20% of the true composition in North skew 2, *Squalus acanthias* had zero observed sequence reads in all three technical replicates. Because these data are compositional, inaccurate estimates of one species can substantially degrade the accuracy of estimates for all species in the community (Fig. 6B). A consequence of getting a single species badly wrong can be seen in the large Aitichson distances indicating low similarity between the true and predicted community (compare y-axes of Figs. 6C, 6D). We present an analysis removing *Squalus acanthias* from the community in Supplement A.

#### Killer Whale Diet

We found modest variation in amplification efficiency among the 10 species in the mock community – notably herring (*Clupea pallasii*) were under-amplified and Atlantic salmon (*Salmo salar*) were over-amplified – but the remaining species were quite similar to one another (Fig. 7). Furthermore, only five of the ten focal species had more than 1% of the reads in any individual sample (Fig. 7A) and herring and Atlantic salmon were not among those five species. As a result, the model had a small (but non-zero) effect on estimated diet composition from both individual samples (Fig. 7A) and the among-sample average diet (7B). Species that were estimated to be above-average amplifiers decreased slightly in the across sample average (e.g. Chinook salmon decreased from 0.531[0.484,0.563] (mean[95%CI]) to 0.512[0.468,0.551] after adjusting for amplification variability) while below-average amplifiers increased slightly (e.g coho salmon increased from 0.225[0.190,0.26] to 0.25[0.198,0.30]). SRKW diet estimates from metabarcoding underscore that when there is little amplification bias among focal taxa, the model output should look very similar to estimates from summarizing raw reads.

**Figure 7:**
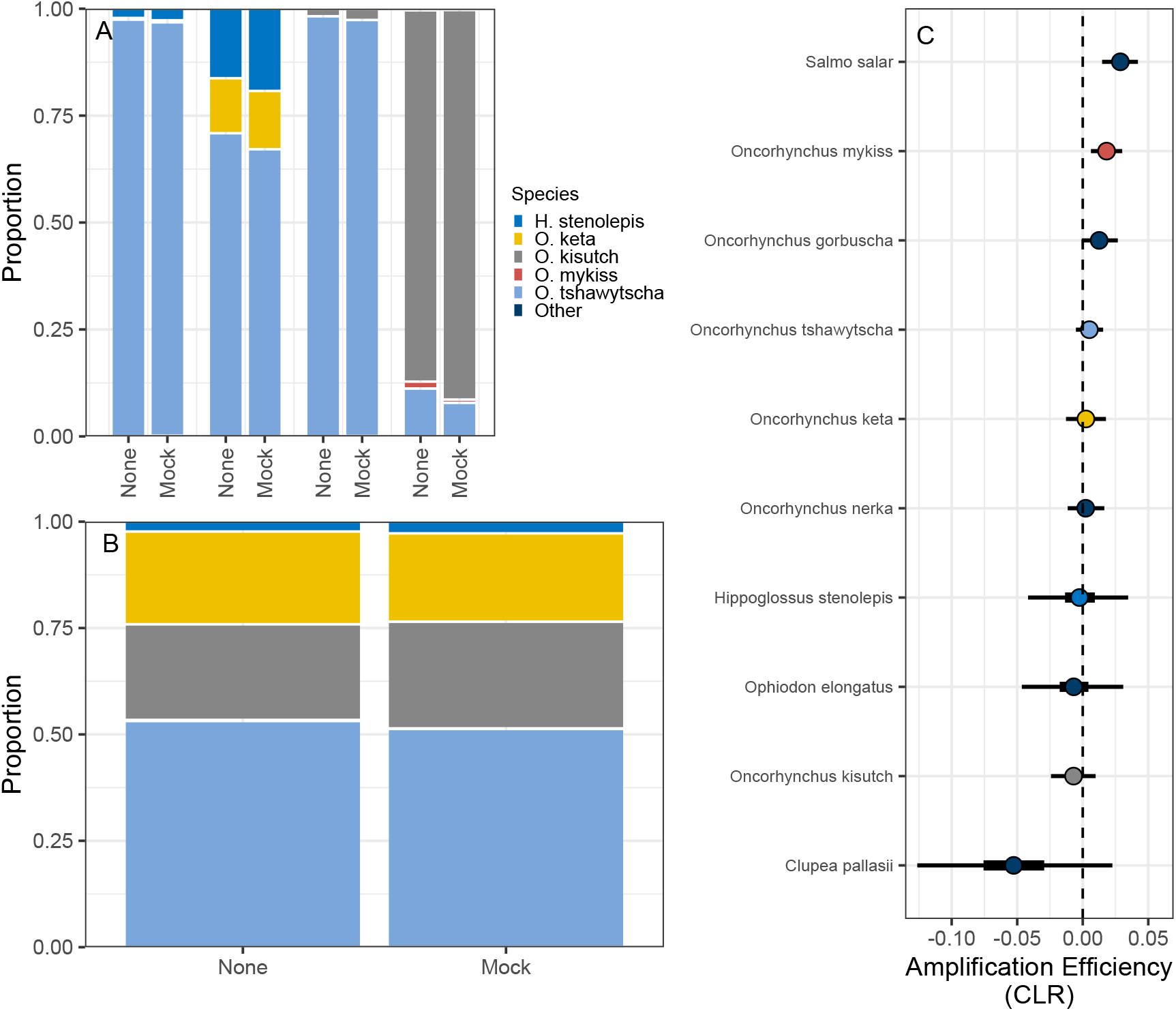
Comparison of calibration methods for southern resident killer whale diets. A: Posterior mean estimates of diet for four individual samples without estimated amplification variability (None) or with amplification variability estimated using a mock community (Mock). B: Posterior mean estimates of diet composition across 8 field samples collected during September betwen 2017 and 2021. C: Posterior mean estimates (interquartile and 95% CI) of relative amplification efficiency after centered log-ratio transformation. Dashed line indicates the geometric mean amplification efficiency among species. Colors correspond to the colors in panels A and B.

## Discussion

By modeling the processes of PCR and sequencing, we provide a general method for correcting for the data distortions generated by amplification bias. The challenges for amplicon data outlined above are general and apply to any kind of multi-taxon PCR-based data. By linking this interpretable, mechanistic model to the existing statistical literature on compositional data analysis, we hope to popularize this technique, such that ecological inferences from metabarcoding may rest on a solid foundation. We find that our approach works well under a wide range of simulated scenarios, and for a diverse set of empirical examples spanning a range of ecological communities and sub-disciplines.

Calibrating metabarcoding datasets is tractable and yields estimates of community composition under many real-world conditions. Importantly, in all of the cases we have examined, accounting for amplification bias improves the estimation of ecological communities relative to approaches that treat raw sequence counts as reflecting underlying communities. However, the approach is not without limitations or uncertainty; where an assay amplifies target species poorly, or where the variance among replicated samples is high – that is, cases where the signal:noise ratio is very low – any inference about ecological communities will be difficult and uncertain.

Thus metabarcoding datasets are very similar to “traditional” (non-molecular) ecological information. Rather than a free-for-all of information having an unknown relationship to the living elements of sampled ecosystems, metabarcoding instead joins visual surveys, nets, traps, culture, and other observation methods in requiring contextual information to become interpretable. Just as cryptic species may elude visual surveys – and so, go unobserved – species that amplify poorly will be rarely observed in the metabarcoding data. Just as it is difficult to predict the abundance of patchily distributed species in a net or visual survey, it is difficult to predict the abundance of a species’ amplicons where the variation among technical replicates is high. And just as researchers should understand the sampling characteristics of their traditional ecological sampling tools in order to best understand the resulting data, so too should researchers understand the behavior of a given primer set in the context of molecular ecological data.

### Calibration, Model Performance, and Extensions

It is now clear that simple tabulations of proportions of amplicon sequence-reads are likely to provide misleading inferences due amplification bias. The model we present corrects for those biases to yield estimates of proportional contributions of each of the taxa (or ASVs, etc) to the original biological sample prior to PCR. Such calibration requires adding information beyond the raw observations of sequence-reads and is not a trivial exercise. However, by dedicating a small portion of a sequencing run to calibration samples, researchers can derive robust estimates of their samples’ underlying DNA compositions. We note that there are several additional paths for calibrating metabarcoding data that are described elsewhere, and these either complement or may be combined with the approaches we discuss (Gold et al. In Review, Hoshino et al. 2021, Silverman et al. 2021).

Our model yields good estimates of community composition, particularly when amplification efficiencies do not vary excessively among taxa and among-replicate variability is low. While simulations suggest that the approach works well for an arbitrarily large number of taxa, in practice, the number of taxa will likely be constrained by the feasibility and patience required to construct mock communities. For valid inference, all taxa of interest must be included and observed in at least one mock community. However, the approach does allow researchers to subset data in arbitrary ways, focusing on only the taxa or samples of interest; model output will reflect this subsetting by estimating the composition of the selected ingroup. Such subsetting already occurs implicitly in most metabarcoding datasets and is evidenced by the species reported to be reliably detected by a given primer and molecular protocol. Therefore measuring and documenting amplification bias is a way of making explicit which components of an ecological community can be measured by given primer and protocol and which cannot.

Intuitively, the model fails in situations in which amplicons provide little information about the original community composition, either because of poor amplification or high variance. Less intuitively, the model can also fail where a few taxa amplify poorly and the rest amplify well – poor estimates for some taxa will affect the estimates for all other taxa because the data are compositional. Iteratively fitting the model can solve this problem, by dropping taxa with low information value and focusing on the remainder. Again, we view this iterative procedure positively because it forces researchers to be explicit about which taxa can be validly assessed by a given metabarcoding approach.

From a statistical perspective, we have presented a relatively simple model and applied it to relatively small datasets. However, the form of the model is easy to extend. The broad suite of statistical tools developed for regression can be easily incorporated; this includes making compositions a linear or non-linear function of covariates, adding random effects, and incorporating spatio-temporal statistical models (see also Supplement A). Furthermore, there are clear paths to generalize this framework to accommodate more than one genetic locus or dataset as well, offering a way of synthesizing information across genetic markers by treating data from each locus as an observation of a common ecological community composition. While such models can be computationally difficult, there are few conceptual blocks to such advancement.

### Practical Problems in Metabarcoding Studies

Our approach links the rapidly expanding field of metabarcoding in ecological applications with statistical methods developed in related fields and offers solutions to several practical problems of amplicon-sequencing techniques. We provide an extensive discussion of practical considerations in Supplement B and focus on a few points of general interest here including decontamination/denoising and the application of traditional ecological statistics to these data.

### Decontamination and Denoising

Whether due to low-level lab contamination (Leonard et al. 2007), index-hopping and related technical problems (Schnell et al. 2015, Costello et al. 2018, Carøe and Bohmann 2020), true detections of unintended taxonomic targets, or other mechanisms, metabarcoding datasets frequently contain observations of non-target taxa. The question then arises whether, or how, these non-target taxa might be responsibly identified and excluded from downstream analysis. For example, in diet-data analysis, might sequences from the host species be safely ignored? Where detections of pigs, chickens, or others arise from PCR reagents themselves, how might we exclude these reads as contaminants?

The technique we present here can be applied to subsets of data by simply excluding non-target taxa, thus changing the denominator for the overall read-depth and shrinking the universe of taxa with DNA proportions to be estimated. Because the model is explicitly compositional, the analysis of any subset of the data remains valid; the resulting estimated proportions will sum to one, reflecting the proportions of the taxa in the subset analyzed, rather than in the entire raw dataset. If, for example, we wish to focus only on the five mammal species present in an environmental dataset, we may analyze only the reads of those five taxa in a set of (say) water samples. The resulting proportions will sum to one, reflecting the contributions of each of the five species to the analyzed subset, saying nothing about the proportions of those five species in the dataset as a whole. We emphasize that this subsetting procedure merely makes explicit what is inherent in any amplicon study: the amplified molecules observed reflect some, but not all, of the molecules in the environment, namely those templates susceptible to the assay being used.

### Diversity Indices and Existing Ecological Statistics

Ecological studies frequently use Shannon, Simpson’s, or other summary indices of diversity. However, sequence reads – whether in raw form or monotonically transformed – do not lend themselves to such indices, given that the indices rely upon proportional estimates of the underlying species present. For example, Shannon Entropy 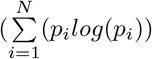, for proportions *p* of species *i* = 1, …, *N*), when applied to raw metabarcoding reads, is meaningless and divorced from its connection to the underlying biology. In general, we care not about *p* as the probability of observing a sequence read from a given taxon, but rather *p* the probability of observing evidence of the underlying species itself, prior to distortion by PCR. Our model explicitly estimates the proportion of species’ DNA in the sample, and so its output is appropriate for these common diversity indices and other standard downstream ecological analysis appropriate to proportion data. We note, however, the proportion of species DNA present does not necessarily reflect the proportion of species counts or biomass present in the environment due to generating, decay, and fate-and-transport phenomena (Barnes and Turner 2016).

### The Need for Replication

Technical replication supplies the data necessary to evaluate the signal:noise ratio in any study. Where replicates are available, our model treats overdispersion – additional variance relative to that expected under a multinomial sampling model – as a random variable (_ϵ*i*_) at the level of biological samples, drawn from a common distribution having a standard deviation *τ*. Replication is expensive in metabarcoding studies due to sequencing costs and labor. Thus it is often desirable in practice to minimize the amount of technical replication in a study, in order to maximize effort elsewhere. In the absence of technical replicates, strong assumptions must be made about the amount of variability in observed sequences. The example of British Lakes (Fig. 5) presents a potential consequence of no replication – overly precise estimates of community composition arise from assuming a multinomial likelihood. However, one might minimize replication after developing enough familiarity with a system to confidently assert a parameter value for over dispersion. Another approach might be to replicate some – but not all – samples in a study, yielding estimates of overdispersion that can be used throughout the dataset. Understanding the origin of overdispersion in sequence counts and best modeling approaches to account for such effects is a significant area deserving of further research in the metabarcoding community, perhaps most especially as to handling zero counts arising from a variety of mechanisms (Egozcue et al. 2020, Silverman et al. 2020).

## Conclusion

Across a broad swath of ecological, microbiological, and biomedical studies, it has become clear that simple read proportions or monotonic transformations calculated from metabarcoding studies have the potential to be deeply misleading. We outline approaches to correct for biases introduced by metabarcoding processes, but acknowledge that the laboratory and statistical effort to adjust for these biases are non-trivial and will hinder their rapid adoption. However, in such a rapidly advancing field, we trust that our work will lead to improved laboratory methods and statistical sortware to make implementation of these approaches routine.

## Acknowledgements

We thank M. Fisher, J. Samhouri, K. Vennemann, and O. Wangensteen for constructive comments on previous versions of the manuscript. We thank B. Hänfling and coauthors for providing excellent data and supplementary materials in their 2016 paper.

## Notes

### Competing Interest Statement

The authors have declared no competing interest.

