## Supplement A for "Toward quantitative metabarcoding"

### Supplement A: Model extensions

Shelton et al

**References** **13**

### Model details

#### Prior Distributions and Model Diagnostics

An important aspect of Bayesian analysis is specifying prior distributions. Table A1 provides prior distributions for estimated parameters in all presented models. All prior distributions are diffuse and independent.

| Parameter | Prior Distribution |
|---|---|
| $\alpha$ | $Normal(0, 0.1)$ |
| $\beta$ | $Normal(0, 10)$ |
| $\tau$ | $Gamma(1.5, 1.5)$ |

Table A1: Prior distributions used in all models

We implemented all models in *R* (v.4.1.2, R Core Team 2021) using the *R* implementation of the Stan language (*Rstan* v.2.21.2, Stan Development Team 2020). For all models we ran multiple chains (British lakes models: 5 chains, 500 warmup, 1000 sampling iterations; Pacific ocean fish models: 3 chains, 1000 warmup, 1500 sampling iterations; southern resident killer whale models: 3 chains, 1000 warmup, 1500 sampling iterations; bacterial microbiome models: 3 chains, 500 warmup, 1000 sampling iterations). All relevant code and data for implementing the model are provided in the accompanying data repository. We used traceplots and $\hat{R}$ diagnostics to confirm convergence ($\hat{R} < 1.01$ for all parameters). There were no divergent transitions in any of the sampling iterations.

### Supplementary Analyses

#### Mock Communities

There are several complementary ways to display the divergence between the true composition of mock communities and the community composition based on simple summaries of the observed reads. One informative way is to provide a way to determine if particular species are persistently under or over represented across multiple mock communities. Here we replot the data presented in Fig. 1 of the main text but rather than highlighting a single community, we focus on a few species and examine those species across several communities (Fig. A1). Some species are repeatedly over-represented in the sequences observed relative to their true abundance (e.g., *G. cernua* in Fig. A1B, *O. elongatus* in Fig. A1D) while others are underrepresented in the observed sequences (e.g., *T. tinca* in Fig. A1B, *M. productus* in Fig. A1D) and still others are near their true proportion.

Left panels show the observed and true proportions

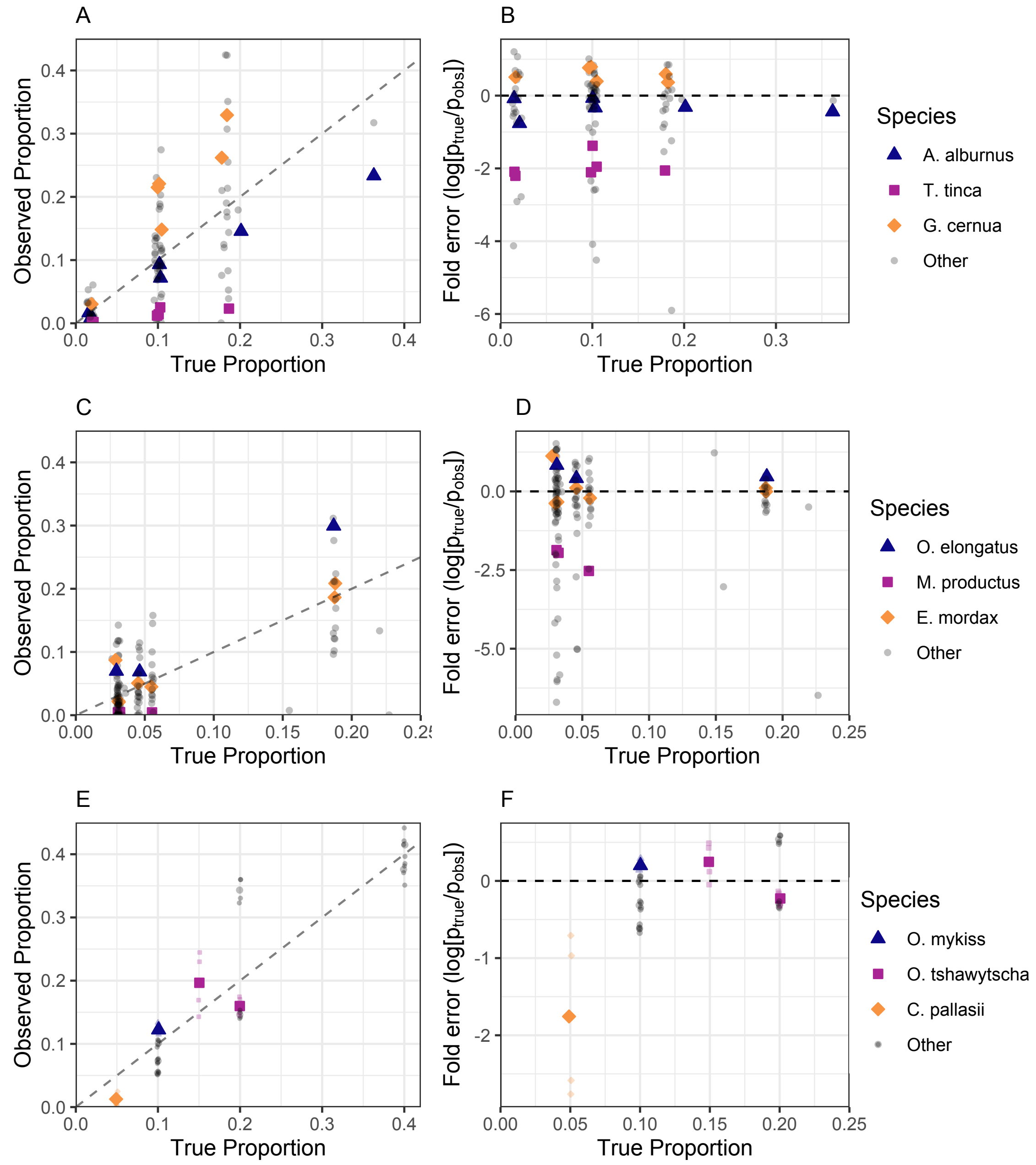

Figure A1: Comparison mock communities presented in Fig.1 of the main text but with individual species highlighted. Rows show results for species within mock communities constructed by Hänfling et al. (*A, B*: ten mock communities of freshwater fishes; cytochrome b mtDNA), and for this study (*C, D*: mock communities of Pacific Ocean fishes; *12S* rRNA; and *E, F*; mock communities of Southern-Resident Killer Whale fecal samples; *16S* rRNA). *A, B, C* compare the observed and true proportions while panels *D, E, F* show fold-error estimates for each community. Each point represents a single species in a mock community. For *A*, each point is a single technical replicate while in *C* and *E*, each point is the average across three technical replicates. Fold-error for each individual species in each community is shown in *B, D, and F*.

#### British Lakes

For completeness, we present out-of-sample predictions for all 10 mock communities presented in Hänfling et al. (2016) (Fig. A2). As described in the main text, two different models are presented here fit. In one, the odd-numbered communities were used as mock communities and the even communities were predicted for use it out-of-sample cross validation. In the other even numbered communities were used as mock communities and the odd numbered communities were predicted for use in out-of-sample cross validation.

We show a measure of predictive accuracy (Aitchison distance) for all 10 communities in Fig. A3. Note that community MC07 is a notable outlier, with a much larger difference between prediction and true composition than any other species. This is largely driven by a single species (*Gobio gobio*) in community MC07 that had zero observed sequences in the data but in truth composed slightly more than 1.8% of the mock community (see also Fig. A2). All other species were closer to their true composition after mock community calibration, but Aitichson distance is almost entirely driven by a single species. This result emphasizes the importance and leverage of zero observations in metabarcoding data (McLaren et al. 2019, Silverman et al. 2021) and in the analysis of compositional data in general (Aitchison 1986).

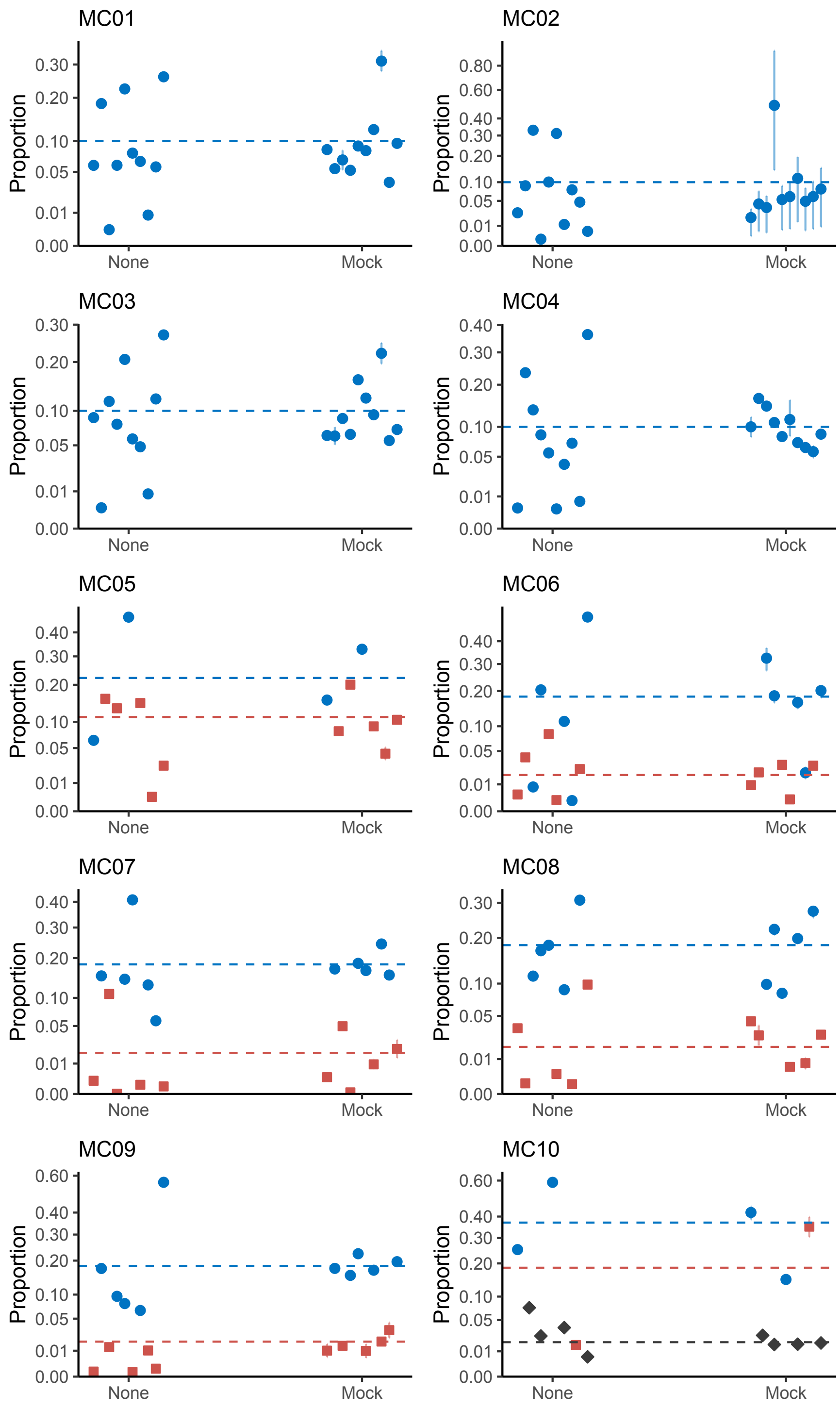

Figure A2: Comparison of calibration methods for British Lakes fish communites. Each panel shows a distinct mock community (MC01 - MC10). Points and error bars show posterior mean estimates (95% CI). Dashed lines shows the true composition for each species with color indicating the true composition for each point. Estimates from a model without amplification variability ('None') are shown alongside estimates from models including amplification variability ('Mock').

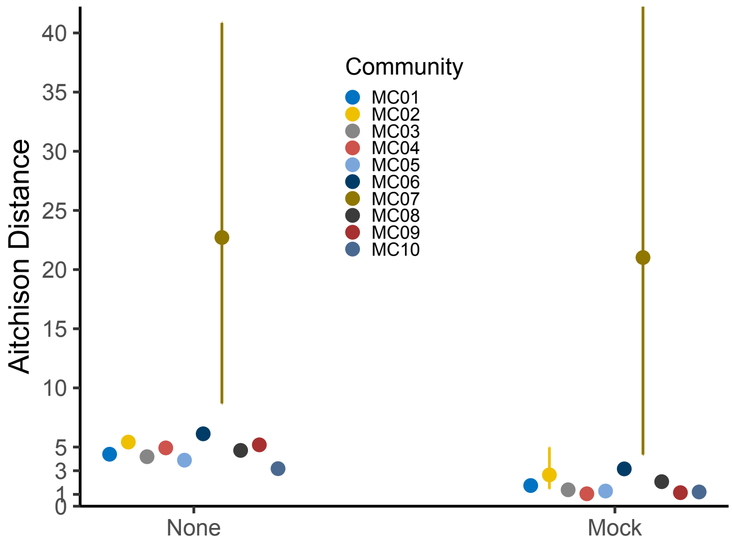

Figure A3: Comparison of calibration methods for British Lakes fish communites. Each panel shows a distinct mock community (MC01 - MC10). Points and error bars show posterior mean estimates (95% CI). Dashed lines shows the true composition for each species with color indicating the true composition for each point. Estimates from a model without amplification variability ('None') are shown alongside estimates from models including amplification variability ('Mock').

#### Pacific Ocean Fish.

For completeness, we present all four communities (Ocean skew 1 and 2, North skew 1 and 2) used for out-of-sample cross validation for the Pacific ocean fish community (Fig. A4). The Ocean skew 1 (Fig. A4A) and North skew 2 (Fig. A4D) communities are shown in the main text (Fig. 6). In all cases, using mock community calibration leads to higher similarity between the true and predicted community composition (Fig. 6).

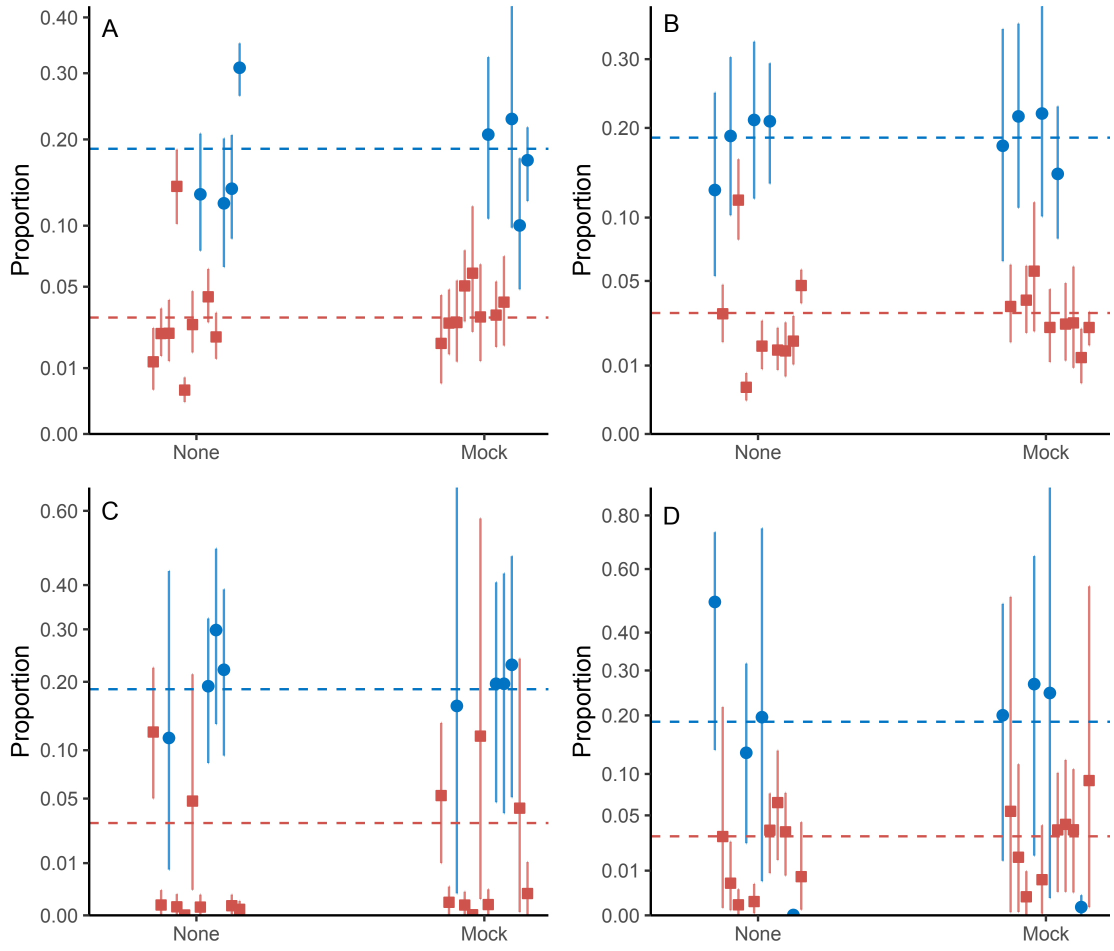

Figure A4: Comparison of calibration methods for Pacific ocean fish communites. *A*: Posterior mean estimates (95% CI) of species composition for the 12 species in the mock community Ocean skew 1 without estimated amplification variability ('None') or with amplification variability estimated using a mock community ('Mock'). Dashed blue line shows the true composition for the species identified with a circle. Dashed red line shows the true composition for the species identified with a square. *B*: Posterior mean estimates (95% CI) of species composition for the 12 species in the Ocean skew 2 mock community. *C* Posterior mean estimates (95% CI) of species composition for the 12 species in the mock community, North skew 1. *D* Posterior mean estimates (95% CI) of species composition for the 12 species in the mock community, North skew 2.

**Subsetting communities.**

To illustrate the value of being able to select a subset of species for analysis, we reanalyzed the North mock communities using North even as a mock community and North skew 1 and North skew 2 as predicted communities. Specifically, we did not reestimate the model, but we did exclude the poorest amplifying species (*Squalus acanthias*, Fig. 6) from predictions (i.e., we made predicted species compositions for all species except *S. acanthias*). Excluding *S. acanthias* changes the true proportions for the remaining species (compare the y-axis in Fig. A5; *S. acanthias* originally comprised 3.125% of North skew 1 and 18.75% of the North skew 2; Table C2). Predicted species compositions after removing *S. acanthias* using mock calibration were superior to using no calibration (North skew 1: Aitchison distance of 10.6 and 12.1 for mock calibaration and no calibration, respectively; North skew 2: 5.3 and 3.6 for mock and no calibration, respectively). For both North skew communities, the Aitchison distance is much smaller for the community excluding *Squalus acanthias* than for the full communities (see Fig. 6) indicating that a single species was largely driving differences between the true and predicted compositions.

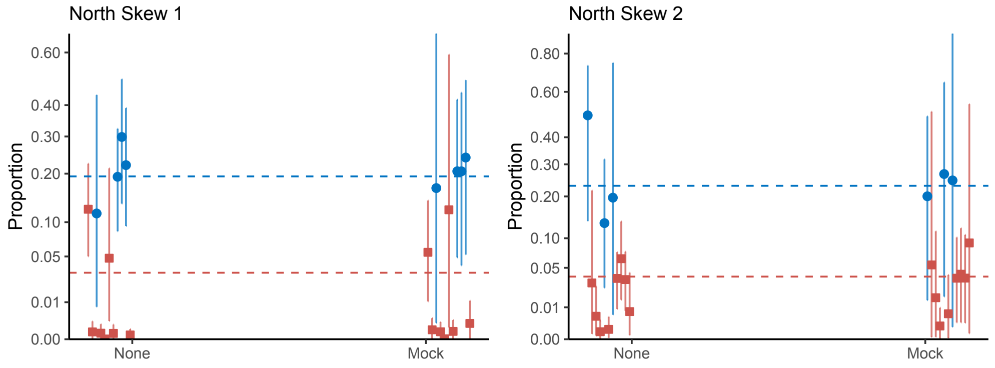

Figure A5: Comparison of calibration methods for the North skew 1 and North skew 2 Pacific ocean fish communites excluding *Squalus acanthias*. Posterior mean estimates (95% CI) of species composition for the 11 species in the mock community without estimated amplification variability ('None') or with amplification variability estimated using a mock community ('Mock'). Dashed blue lines show the true composition for the species identified with circles. Dashed red lines show the true composition for the species identified with a square.

#### Southern Resident Killer Whales

For completeness, we present the estimated composition of the eight field samples of southern resident killer (SRKW) whale diets (Fig. A6). Of note is that for any individual sample, relatively few species are detected. Of the eight samples, four were estimated to have a majority of Chinook salmon *Oncorhynchus tshawytscha*, two have a majority of chum salmon, *O. keta*, and two have a majority of coho salmon, *O. kisutch*. Because the amount of change due to amplification bias depends on the identity of species detected in each sample, and their relative observed abundances, calibrating for amplification bias results in substantial shifts in species composition in some cases and small changes to species composition in other cases. It is important to remember that these samples are not representative of SRKW diet overall, but rather a group of samples that can be used to illustrate the application of the methods to diet data.

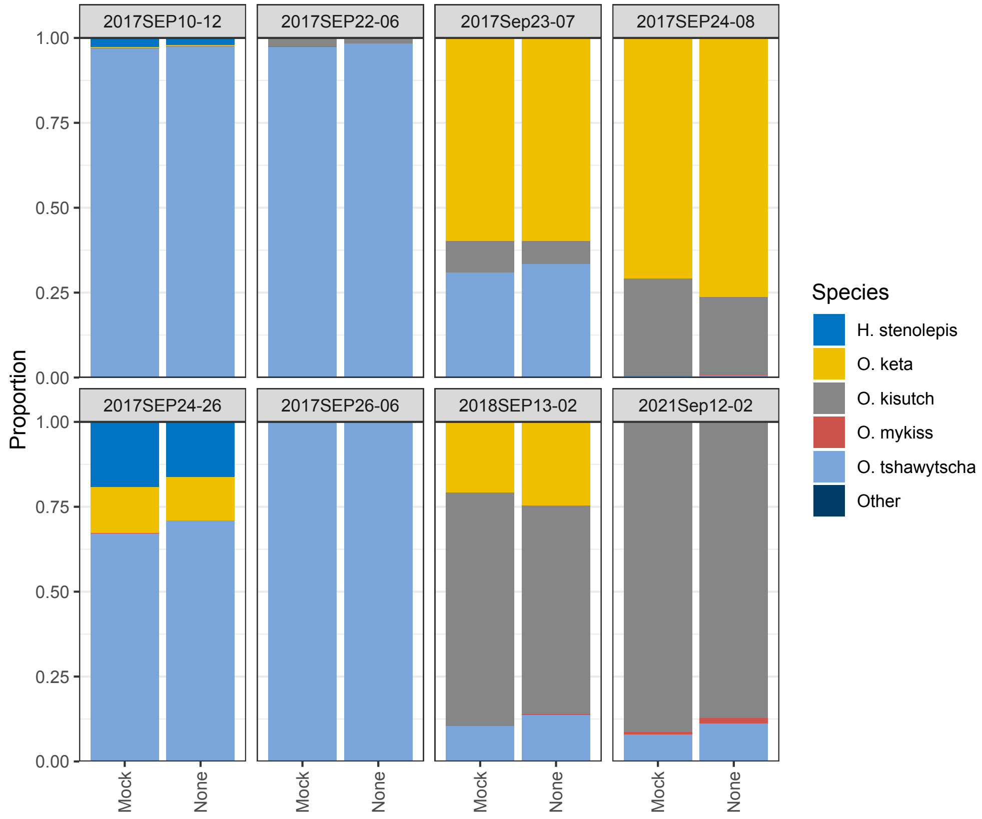

Figure A6: Comparison of calibration methods for Southern Resident Killer Whale diets in eight individual samples. Each facet shows the posterior mean estimate of diet for eight individual samples without estimated amplification variability ('None') or with amplification variability estimated using a mock community ('Mock').

#### Bacterial Microbiome Data

We extracted data from Gohl et al. (2016) to apply our approach to metabarcoding data derived from bacterial microbiome communities. We focused on data from two communities consisting of 17 taxa (Table C5). There were three technical replicates for each community. We used the "even" community as our known mock community and predicted the "staggered" mock community (Table C5). We then compare estimates from our model and the proportions estimated from raw reads against the true proportions (Fig. A7). While proportions derived raw reads often deviated substantially from the true proportion, predictions from the model calibrated with the even community matched the true proportions closely. For 16 of the 17 species the 95% prediction interval included the true estimate with only *Acinetobacter* having a true value slightly outside of the 95% interval (Fig. A7).

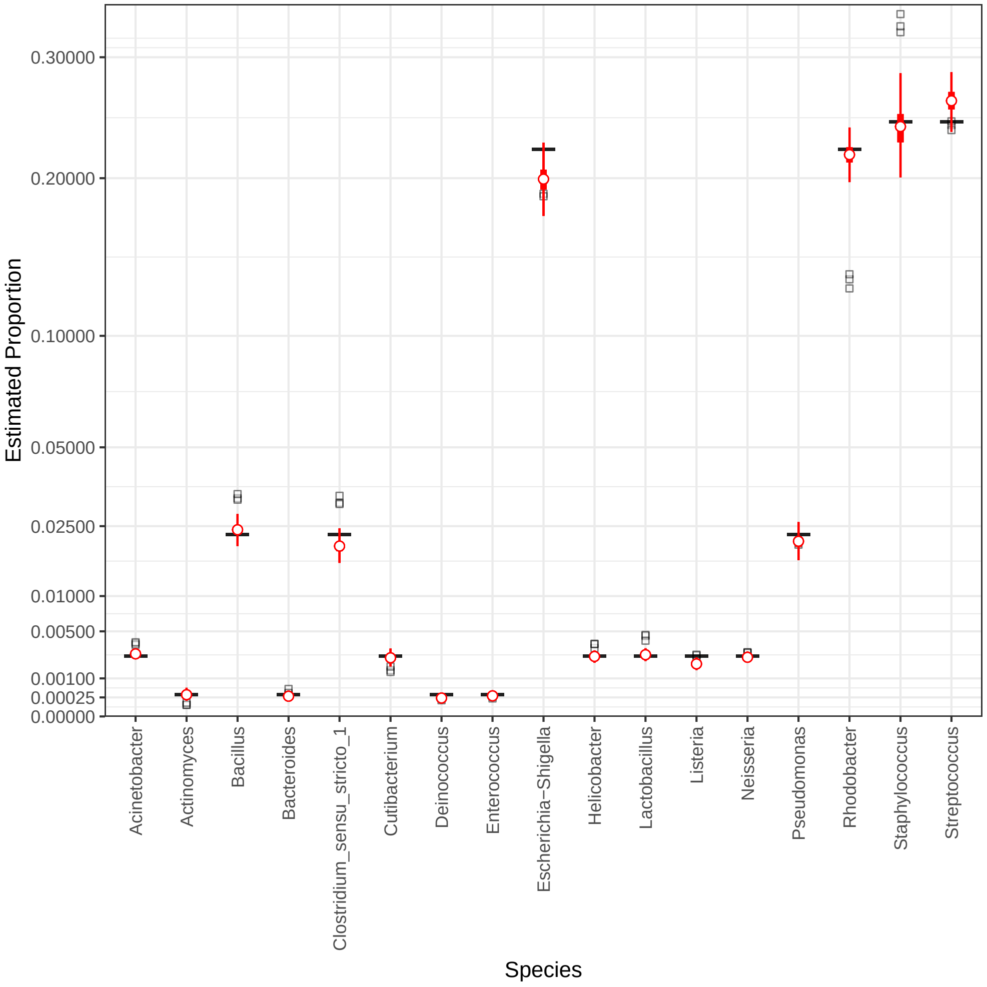

Figure A7: Predictions for the 'staggered' microbiome community. Black horizontal line indicates true proportion, squares show estimates from individual replicates, red points and lines show posterior mean, interquartile range, and 95% CI.

### Model extensions and alternative parameterizations

#### Model Extensions

In the main text we present a straightforward statistical model with easy to interpret notation. In this section we outline immediate extentions to the main text model to make clear the connection of our work to the broader field of generalized regression models as well as note connections to the broader world of compositional analysis. To reiterate a point from the main text, models for compositions have been extensively investigated in other fields, from geology (Buccianti et al. 2006) to habitat selection (Aebischer et al. 1993). Researchers using metabarcoding data can learn from the extensive experience using these kinds of models in other fields to rapidly advance their use. The most straightforward extention is to illustrate how to include samples associated with specific sampling sites (indexed by $j$, $j = 1, 2, ..., J$) and technical replicate ($k$, $k = 1, 2, ..., K$) can be included in the model. For notational simplicity we assume there are a total of $JK$ observations associated with unique site-technical replicate combinations. We can modify eq. 6 from the main text to reflect sites and let $\boldsymbol{X}$ be a design matrix containing covariates (with $C$ columns and $JK$ rows; covariates can be factors or continuous or a mix), $\boldsymbol{\beta}_i$ be a vector of coefficients for for length $C$ for each $i$ species. Then the full model can be written

$$\boldsymbol{Y}_{jk} \sim Multinomial(\boldsymbol{\mu}_{jk}, N_{jk}) \tag{A.1}$$

$$\mu_{ijk} = \frac{e^{\nu_{ijk}}}{\sum_{i=1}^{I} e^{\nu_{ijk}}} \tag{A.2}$$

$$\boldsymbol{\nu}_i = \boldsymbol{X}\boldsymbol{\beta}_i + N_{PCR}\alpha_i + \boldsymbol{\epsilon}_i \tag{A.3}$$

$$\epsilon_{ijk} \sim N(0, \tau_i) \tag{A.4}$$

Then composition of each species after correcting for amplification variability ($N_{PCR}\alpha_i$) and overdispersion ($\boldsymbol{\epsilon}_i$) is defined by $\boldsymbol{X}\boldsymbol{\beta}_i$. Specifically, the predicted proportion for species $i$ in site $j$, $\pi_{ijk}$, would be

$$\boldsymbol{\eta}_i = \boldsymbol{X}\boldsymbol{\beta}_i \tag{A.5}$$

$$\pi_{ijk} = \frac{e^{\eta_{ijk}}}{\sum_{i=1}^{I} e^{\eta_{ijk}}} \tag{A.6}$$

and all technical replicates at a site would have identical predictions.

While this model simply adds fixed covariates to the model, there is no reason that the general regression form above cannot contain additional elaborations (e.g., non-linear effects or spatio-temporal effects). Silverman et al. (2021) discuss parameterizations that are appropriate for efficient computation of this type of latent variable models.

#### Alternative parameterizations

In discussions about this topic, experts on metabarcoding data have mentioned that the process model for PCR outlined in eq. 1 is not accurate. Specifically, they objected to the notion that the amplification efficiency is constant across PCR cycles and posited that the efficiency at the first few cycles could differ substantially from later cycles. While there is some empirical evidence for consistency in amplification across PCR cycles (e.g., Silverman et al. 2021), it is not overwhelming. Therefore it is reasonable to consider alternate parameterizations of our model that do not require a constant amplification efficiency.

Let us consider the general case where each PCR cycle has a distinct amplification efficiency so across the entire PCR reaction, the expected number of amplicons is a product of the amplification efficiency at each cycle,

$$A_i = c_i \prod_{k=1}^{N_{PCR}} (1 + a_{ik}) \tag{A.7}$$

where $a_{ik}$ is the amplification efficiency for taxon $i$ at cycle $k$ in a given protocol. As we did with the model in the main text, we can take ratios between species and logarithms. For notational simplicity, let $g_i = \prod_{k=1}^{N_{PCR}} (1 + a_{ik})$, then,

$$\log\left(\frac{Y_i}{Y_j}\right) \propto \log\left(\frac{c_i}{c_j}\right) + \log\left(\frac{g_i}{g_j}\right) \tag{A.8}$$

where we include observations for two species $i$ and $j$. As in the main text, we can arbitrarily define one taxon to be a reference taxon ($R$) and define a new set of parameters relative to this reference taxon. Let $\beta_R = 0$ be the log-abundance of the reference taxon in the initial sample and $\beta_i$ be the abundance of taxon $i$ relative to the reference ($\beta_i > 0$ indicates taxon $i$ is more abundant than the reference, $\beta_i < 0$ the opposite). Similarly, let $\gamma_i$ be the log-efficiency over the entire PCR relative to the reference taxon ($\gamma_R = 0$). As before, $\nu_i$ is the log-abundance of taxon $i$ relative to the reference taxon after sequencing,

$$\nu_i = \beta_i + \gamma_i \tag{A.9}$$

Comparing this equation with eq. 3 in the main text, it becomes clear that assuming variable amplification rates results in a functionally equivalent statistical model to that presented in the main text (i.e., $\gamma_i = N_{PCR}\alpha_i$). While $\alpha_i$ defines the relative amplification per PCR cycle, $\gamma_i$ is the relative amplification across all PCR cycles and so these terms will have different magnitudes. In the context of calibration using mock communities, however, writing the model in terms of $\alpha$ or $\gamma$ will have no practical consequence.

Using amplification efficiencies across the entire PCR reaction is the approach adopted and generalized by McLaren et al. (2019) for microbiome data. We suggest interested parties read their work in full.

Stan Development Team. 2020. RStan: The R interface to Stan.
