## Supplement B for "Toward quantitative metabarcoding"

Supplement B: A Practical Guide to Quantitative Metabarcoding

**Frequently Asked Questions about eDNA and Quantitative Metabarcoding and how these relate to the presented model.**

**Q:** Your model of PCR is wrong.

**A:** First, this isn’t a question. But also: no model is perfectly correct. Here, we focus on modeling the process of PCR to determine the varying amplification efficiencies of different taxa that cause the disconnect between the relative proportion of starting DNA concentrations and the relative proportions of amplicons. The model presented here is a simplification of the process, but we demonstrate using both simulated and empirical data that the model performs well. We also show how an alternative parameterization that allows for amplification variability among PCR cycles leads to a very similar statistical model in Supplement A.

We look forward to future iterations that better model the process. For example, the subsampling process in which some molecules from the overall DNA extract in the tube are selected to be template molecules for the PCR can lead to species dropping out stochastically among triplicate reactions, simply because they didn’t happen to be in the (say) 1uL template you grabbed out of the 100uL in the tube. This subsampling creates a special kind of overdispersion in the resulting amplicon counts, where (for a given species) we observe zero reads in some replicates and potentially high read-counts in other replicates (see, e.g., Egozcue et al. 2020). Future work better characterizing such overdispersion, and the occurrence and frequency of zero observations in particular, will aid in model performance.

**Q:** What is the list of important assumptions that go into the statistical model?

**A:** We assume the following:

- The PCR reaction was still in the exponential phase when it was stopped and amplicons were put onto the sequencer.
- The amplification efficiency of the PCR reaction is characteristic of a given taxon-primer combination, and is independent of the other taxa in the template mix.
- Mock communities provide a good estimate of amplification variability for field collected samples of unknown composition.
- The taxon-specific amplicons observed via sequencing are a stochastic sample of the amplicons present after amplification and can be approximated using a multinomial distribution.
- There is additional variability that arises as a result of (a) subsampling the extracted DNA into triplicate technical replicates and (b) PCR amplification. This produces variability in taxon-specific observations among triplicates. This variability can be adequately captured using a random effect for each taxon-sample-technical replicate combination
- We assume all unique ASVs in a sample map to a single taxon and share a single amplification rate. Thus, amplification variability is a taxon-specific process.

**Q:** How do we know if the PCR reaction is still in its exponential phase?

**A:** The assumption for the model that the PCR reaction is still in its exponential phase requires sequencing the same sample several times in order to visualize the change in amplicons at different cycle numbers.

To address this, we conducted a series of lab experiments to confirm that generated mock communities were not saturated. First, we created a mock community by amplifying a ~600 bp fragment of the 12S mtDNA gene from genomic DNA extracted from tissue samples (see Supplement C for more details). Importantly, this allowed us to convert the starting concentration of the mock communities from [ng/ul] as measured by Qubit to [copies/ul] before amplifying the ~170 bp fragment of interest (i.e., MiFish) that was nested within the ~600 bp fragment. Furthermore, after the MiFish PCR, the same conversion from [ng/ul] as measured by Qubit of the amplicons can be converted to [copy/ul] with the adjusted size of the fragment (~170 bp after PCR versus ~600 bp before PCR). The mock community had a concentration of 1.42x10^9 copies/µL before the MiFish PCR amplification. We conducted a 1:10 serial dilution to 1.42 copies/µL and then ran the MiFish PCR with a total of 30 cycles (see supplemental methods) across all serial dilutions and negative control. We then measured the resulting DNA concentration after PCR by Qubit.

If the MiFish PCR had saturated, we would have expected to see a saturation of the total resulting amplicon DNA concentration as the reagents are exhausted. We did not observe evidence of saturation even at 1.42x10^9 copies/µL at 30 cycles (Figure B1). However, in an abundance of caution we chose to use 142 copies/µL as our initial input for Pacific mock communities ensuring that we avoided saturation. We note that this starting concentration is substantially higher than the typical environmental DNA concentrations. This result suggests that most metabarcoding approaches are far from saturation and that the model assumption is reasonable.

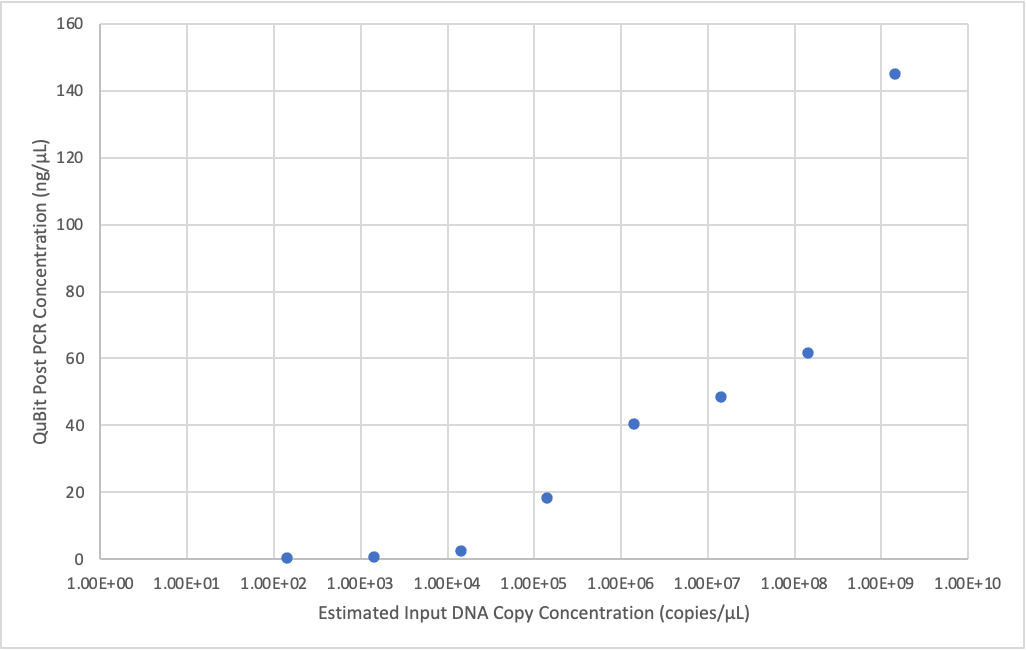

Figure B1: Post-PCR amplification concentrations vs. pre-PCR amplification concentrations of DNA [copies/uL] using serial dilutions of a known mock community.

**Q:** Will saturation in a mock community happen, even if it doesn’t happen in environmental samples?

**A:** We expect different samples to experience saturation at different points in the PCR process, even between environmental samples or mock community samples. The starting concentrations of DNA in the sample, the amount of polymerase and dNTPs, the number of cycles, and other factors will all likely affect PCR saturation. Here we note that the empirical data generated for this paper (see “Pacific Ocean Fishes” in the main text) were mock communities and we determined that the PCR reaction was still in the exponential phase and the model performed well. However, we diluted our mock communities by 8 orders of magnitude to avoid potential PCR saturation. We recommend preliminary testing to check for saturation for both mock communities and environmental samples before applying the model to samples. The primary situation in which saturation is a danger is one in which the PCR reaction is starting with high template concentration and the template molecules have high amplification efficiencies. This is more likely to be the case in mock communities, microbiome, and bulk metabarcoding samples than for environmental samples.

**Q:** The assumption that amplification efficiency is constant and independent of species composition seems a heroic one. Do you have evidence for that?

**A:** The freshwater fish example in the main text, from Hänfling et al. (2016), features independent estimates of the amplification efficiency of each taxon from communities having different community compositions; these sets of estimates are notably similar to one another, evidence that the parameter values are indeed approximately constant. Similarly, the Pacific fish examples in the main manuscript feature estimates of amplification efficiency derived – for each species – from different communities, and the narrow credibility intervals on those estimates again suggest approximate consistency. Likewise, our results from independent estimates of amplification efficiencies from mock communities, joint abundance-metabarcoding models, and variable PCRs support this hypothesis. These different datasets feature different mtDNA loci and appear to behave similarly and thus we feel this is a safe assumption.

**Q:** How does the absolute magnitude of amplification efficiency affect the ability to estimate species compositions?

**A:** When a given primer set amplifies a species very efficiently, we expect PCR signal to be far greater than stochastic noise, and so there should be a lot of real information embodied in the observed amplicon counts for that species. That, in turn, should yield good estimates of starting proportions. But when amplification efficiency is poor – or when a species/template is only very rarely observed, we have little information on which to base an estimate. For those species, we expect poor estimates of starting proportions. Finally, because the data are compositional, poor estimates of some species mean greater uncertainty for the estimates of all other species in a mixture. Accordingly, it may make sense to focus on the species for which the signal to noise is greatest… which is in fact how much of traditional ecological sampling is done (e.g., via visual surveys, focusing on those species that are most easily observable). Elsewhere, we provide an example of such approaches by excluding species from Pacific Ocean mock communities (*Squalus acanthias*) that performed poorly (Supplement A). By removing the species that amplified poorly, we better reconstruct the known input DNA concentrations of the remaining observed species.

**Q:** Is there a cutoff for amplification-efficiency (`a`) values, below which we don’t really trust results?

**A:** The absolute values of `a` are not really important, but the distribution of `a` values among species (within a PCR reaction) is important. In particular, when there is a narrow distribution of values, the model performs better (see Fig. 4 in main text) than when there is a wide distribution of values. We also tested the model by running simulations where most species have a narrow range of `a` values, but one species has a very low value. The model performs well for the species with good `a` values and the single poor `a` values does not perform well. In other words, the one bad apple does not spoil the bunch. However, we suspect that several bad apples do quickly spoil the estimates for a compositional community, because high uncertainty in more than one parameter estimate is likely to ramify throughout the other estimates.

Additionally, we expect that universal metabarcoding primer sets which amplify hundreds of thousands of taxa across dozens of phyla to have lower quantitative capabilities than targeted metabarcoding assays on one class or sub-group because there is higher likelihood that a universal metabarcoding primer set amplifies many taxa poorly. Further experimental and field tests of this hypothesis are warranted, but our results suggest that more targeted metabarcoding assays will result in more accurate quantitative estimates.

**Q:** While we can potentially see the impact of these analyses for providing proportional abundance estimates, we are primarily interested in using metabarcoding for determining the presence or absence of taxa. How does amplification bias affect the determination of presence or absence?

**A:** Unfortunately, determination of a taxon’s presence or absence is affected by amplification bias and the vagaries of compositional data. Just as the abundance of a taxon will be affected by the other species present in a sample, the detection or non-detection of a taxon will be affected by its true abundance, its relative amplification efficiency, and the abundance and amplification of the other species present in that sample. Intuitively, species with higher relative amplification efficiencies and higher abundance will be easier to detect than species with low amplification efficiency and low abundance. As an empirical example, *Squalus acanthias* in our Pacific Fish community (Fig. 6B) was not detected in any of the three technical replicates despite comprising 20% of the community in the North skew 2 community. This occurred because this species amplifies poorly relative to the other species in this mock community. In the North skew 1 community, however, *Squalus acanthias* was observed in two of three technical replicates despite only comprising 3.125% of the initial mock community (Fig. A5).

Overall, the phenomenon of amplification variation means that each taxon will have a distinct and context dependent probability of detection. This result is currently not generally accounted for in occupancy models used for ecological metabarcoding data.

**Q:** Can I use the same amplification estimates for one primer set even if I use different PCR conditions, enzymes, and reagents?

**A:** We think that amplification efficiency of a given species is modified by both the cycling conditions, DNA polymerase, and the reagents used to conduct the PCR. Any changes to the PCR conditions will affect the affinity of the primers to the DNA molecules and then impact the resulting efficiency of the reaction. For example, including a touchdown PCR changes the effective number of PCR cycles that any given molecule will experience; those molecules binding to primers at higher temperatures (i.e., those with more exact nucleotide matches) will then experience a greater number of cycles than will those that bind at lower temperatures. Because molecules from different taxa will be subject to different (and unknown) numbers of cycles, this will change the calculated amplification efficiency for each taxon by an unknown amount. In sum, yes, cycling conditions, enzymes, and reagents of PCR reactions probably do matter. As of writing of this manuscript, we don’t know exactly how stable an amplification efficiency for a given species and primer set is across changes in PCR chemistry and conditions. However, if you are using the same protocol for all samples, it should not affect the model performance. The caveat is that separate independent estimates of amplification efficiencies from mock communities will need to be generated for each unique PCR protocol until further research demonstrates the stability of amplification efficiencies across these parameters.

**Q:** I actually run several PCRs with a subsampling step in the middle. Does that mess up my data or violate model assumptions?

**A:** Subsampling occurs at the beginning of the process by subsampling DNA extract to put into the PCR reaction, and at the end of the process by subsampling amplicons to sequence (see Fig. 2 in the main text). Adding another subsampling process of partially amplified amplicons to then continue to amplify does not necessarily violate the model assumptions, but it likely would add additional variability to the observations. The model captures this variability in the overdispersion term and therefore should be able to account for such subsampling.

**Q:** Mock communities are hard to make - do I have to make them?

**A:** Mock communities are difficult to make because doing so requires tissue samples for every species of interest, which in many cases may be hard to find or is impractical given the sheer number of target taxa. However, we do find that the mock community calibration method is superior to the variable PCR cycling calibration method from our simulation results. We recommend including species of utmost importance in a mock community in order to really understand relative amplification efficiencies of the most important taxa. Data can be subset to include only the species included in the mock community to determine relative starting DNA concentrations of the species of high priority. Other calibration methods like variable PCR cycling may also successfully allow the user to estimate amplification efficiencies and thereby fit the model we describe here as demonstrated in Supplement A.

One suggested approach based on our current results is to combine variable PCR cycling with mock communities to provide multiple independent estimates of amplification efficiencies. For example, if you have 100 environmental samples one could take 10 µL from each sample and make a “super concentrated pool” of all available DNA. However, one pool alone will likely dilute out rarer targets, particularly ones that are only abundant in a subset of the samples. Thus, one could make a series of concentrated pools based on treatments or subsets of the data to avoid dilution of rarer targets. Such determinations can be made after sequencing all samples individually to allow for appropriate coverage of representative samples. In addition, to the variable PCR cycling, mock communities can be made for the subset of important taxa of interest. Together both can provide amplification estimates for a wider range of taxa, while also providing higher quality estimates of key taxa. See Silverman et al. (2021) for an extensive discussion of the variable PCR approach.

**Q:** If I do make mock communities, how many species in the sample community need to be in the mock community? Must it be taxonomically diverse? Should I make more than one mock community?

**A:** The number of species and taxonomic diversity included in your mock communities depends upon your taxa of interest. Highest-priority taxa should be included in your mock community, giving you the most information about their behavior in a metabarcoding study. Our empirical data suggest that even two species within the same genus can have different amplification efficiencies so any species of interest should be included even if they are closely related. Including multiple mock communities with overlapping species or different starting proportions will improve model estimates but is not required.

**Q:** Do I need technical replicates to run the model?

**A:** We highly recommend including technical replicates of PCR reactions, for reasons discussed in the main manuscript. Replication is not required to run the model (see empirical data by Hänfling et al. (2016) in the main text), but without replication there is no way to estimate overdispersion in your observations. We fully recognize that this increases the costs of associated sampling efforts.

**Q:** Can you run more than one primer with this model?

**A:** Not currently, but with some manipulation, yes. A given species will have different amplification efficiencies for each primer set used. Therefore, integrating information from multiple primer sets will improve quantitative estimates of the underlying starting DNA concentrations, but the model is not currently written to process multiple primer sets and integrate the results.

**Q:** Does it really not matter what my reference species is?

**A:** In general, the reference species will not have substantial practical consequences for model performance because all species will be measured relative to the same (arbitrary) choice. However, if by chance the reference species is rarely observed or has a very low true amplification efficiency – and hence, the model has little information on which to base a reference estimate – the model will likely struggle. There are solid arguments for using log-ratio transformations other than the additive log-ratio transform we employ (e.g., centered log-ratio or isometric log-ratio transforms) due to superior geometric and statistical attributes with respect to compositional analysis. However, a detailed discussion of these topics are beyond the scope of this paper. We refer interested readers to entry points to the broader literature on compositional analysis (Aitchison 1986, Pawlowsky-Glahn and Buccianti 2011).

**Q:** I see that you can observe zero amplicons after sequencing. Does observing zero sequences mean that a given species was absent from the sample?

**A:** Understanding and dealing with zero observations is one of the most complex and difficult topics in the analysis of compositions. Silverman et al. (2020) does a good job of describing multiple different processes that can lead to observing zero sequencing reads in sequence data. Our model deals with zero observations via the multinomial likelihood (eq. 5 in main text), but it does not differentiate among the different possible processes that can lead to an observation of zero sequences. Because the model is largely evaluated in log-space, we cannot allow for any taxon to be truly absent from any site (in terms of the model presented in eqs. 5-8, $e^{\nu_{i}}>0$; because the logarithm of zero is undefined). Instead, we assume that all taxa in the model are present in every sample, but occur at a very low level (e.g., taxon X might comprise a proportion of 0.00000001). Thus, we use very small numbers that are essentially zero to apply the required mathematics to run the model. These numbers are determined by the prior distributions for the various parameter in the model, and the prior distribution for $\beta$ in particular (see Supplement A). This is akin to ecologists’ use of log(n+1) in visual surveys or manual counts.

**Q:** What will the difference be if I run the model using species vs. genera or higher taxonomic groupings?

**A:** We know that different species have different amplification efficiencies for a given primer set (Figs. 5, 6, 7). Our results also indicate that the amplification biases between species are for the most part independent of how closely related species are. In other words, it does not seem like amplification efficiency is the same (or even similar) for two species in the same genus or family. Rather, at least at the scale of fishes, amplification efficiency is not closely tied to taxonomy. Therefore, grouping any species – which, again, will amplify differently – into a synthetic unit for analysis will result in far greater uncertainty in the posterior parameter estimates, and is likely to lead to disaster. (Yes, at some phylogenetic scale, amplification efficiency must be associated with relatedness, since the nucleotide sequences in the primer sites are phylogenetically associated. However, within large taxonomic groups, `a` values appear to be largely unrelated to one another.)

**Q:** Might there be intraspecific variation in amplification efficiency?

**A:** Sure. We often have multiple ASVs that assign to the same species, raising the question of whether these different ASVs have different values of `a`. To try to answer this question, we looked at the number of ASVs per species in our ocean empirical dataset in the main text. First, of the 88 unique species detected in our mock communities, most had multiple ASVs: only 12 species (~14%) were represented by a single ASV. The range of number of ASVs per species was from 1 to 287 ASVs, with over half having more than 10 ASVs (mean: 43, median: 6). In the main text, we summed all the reads from all of the ASVs assigning to a single species to run the model. To check the relationship between ASVs within a species, we took only the top ASV per species and removed all other ASVs and ran the model. We found that the model performed very similarly, hardly changing the parameter estimates for species and not changing the rank abundance of alpha values between species. This suggests that although there may be many ASVs per species, the top ASV is driving the amplification efficiency for the species. Thus, at least in our dataset, there might be intraspecific variation in amplification efficiency, but either the differences are very small, or the top ASV within a species drives the estimates.

**Q:** Should I worry about contamination?

**A:** Contamination arises from a few different processes. At each step in the process, contamination can occur. This includes during field sampling, DNA extraction, PCR amplification, bead cleaning amplicons, adding indexes by PCR, and pooling samples before sequencing. Contamination can be the addition of DNA or amplicons from the field or laboratory that are not from the samples (e.g., from reagents or some other species processed in the lab from a different project, or from human). There can also be cross-contamination where DNA or amplicons from one sample get into a different sample. The model does not fix any type of contamination, but in the case that contamination does occur, identified contaminants can be excluded from the dataset and the model can be run while ignoring those species. This is preferable to the common practice of simply subtracting reads from species identified as contaminants due to the compositional nature of sequencing data and the different amplification efficiencies across species (whether they are contaminants or not).

**Q:** Relatedly, can I trim my dataset to focus only on the species I care about?

**A:** Yes, that’s fine. The model results will be proportions of the taxa you are focused on (the taxa remaining in the analysis after trimming).

**Q:** Can I exclude host DNA from my diet study?

**A:** Yep, same answer as above.

**Q**: Can I revisit analyses from previous metabarcoding studies by adding a new post-hoc mock community?

A: Yes. The only caveats are that you want to make the protocols, primers, polymerase and other laboratory methods for the new mock communities as similar as possible to the original data to ensure comparability between the mock community and the field samples.

**Q**: Can I reuse the same amplification efficiency for a species-primer-condition combination from one study to another?

A: We believe this should work, yes. But this deserves additional corroboration and scrutiny.
