## Supplement C for "Toward quantitative metabarcoding"

**Q:** How do we know if the PCR reaction is still in its exponential phase?

**A:** The assumption for the model that the PCR reaction is still in its exponential phase requires sequencing the same sample several times in order to visualize the change in amplicons at different cycle numbers. We conducted a series of lab experiments to confirm that generated mock communities were not saturated during PCR before fitting the model to the data. We conducted an 8-point 1:10 serial dilution to starting concentrations for one of our mock communities and then conducted the 12S MiFish PCR with a total of 30 cycles (see Supplement C) across all serial dilutions and negative control. We measured the resulting DNA concentration after PCR by Qubit and converted from mass back to copy number.

If the PCR reactions had saturated, we would have expected to see a plateau of the total resulting amplicon DNA concentration as the reagents are exhausted. We did not observe evidence of saturation even at our highest dilution (1.42x10^9 copies/µL) at 30 cycles. However, to be conservative and to simulate more realistic concentrations from environmental samples, we chose to use ~100 copies/µL as our initial input for Pacific mock communities. Because environmental samples are likely lower in initial concentration, this suggests that most metabarcoding approaches are far from saturation and that the model assumption is reasonable.
